# An Amygdalar Oscillator Coordinates Cellular and Behavioral Rhythms

**DOI:** 10.1101/2024.03.08.584151

**Authors:** Qiang Liu, Jiali Xiong, Dong Won Kim, Sang Soo Lee, Benjamin J. Bell, Chloe Alexandre, Seth Blackshaw, Alban Latremoliere, Mark N. Wu

## Abstract

Circadian rhythms are generated by the master pacemaker suprachiasmatic nucleus (SCN), in concert with local clocks throughout the body. While many brain regions exhibit cycling clock gene expression, the identity of a discrete extra-SCN brain oscillator that produces rhythmic behavior has remained elusive. Here, we show that an extra-SCN oscillator in the lateral amygdala (LA) is defined by expression of the clock-output molecule mWAKE/ANKFN1. mWAKE is enriched in the anterior/dorsal LA (adLA), and strikingly, selective disruption of clock function or excitatory signaling in adLA^mWAKE^ neurons abolishes Period2 (Per2) rhythms throughout the LA. mWAKE levels rise at night and promote rhythmic excitability of adLA^mWAKE^ neurons by upregulating Ca^2+^-activated K^+^ channel activity specifically at night. adLA^mWAKE^ neurons coordinate rhythmic sensory perception and anxiety in a clock-dependent and WAKE-dependent manner. Together, these data reveal the cellular identity of an extra-SCN brain oscillator and suggest a multi-level hierarchical system organizing molecular and behavioral rhythms.

## INTRODUCTION

Daily rhythms of physiology and behavior in mammals are organized by a master circadian pacemaker in the suprachiasmatic nucleus (SCN)^1–4^. Beyond the SCN, local oscillators throughout the body are thought to promote specific biological rhythms^1,5–8^. For example, work over the past few decades has demonstrated the importance of peripheral oscillators (such as in the liver) in regulating the cycling of metabolic processes^7,9^. In the brain, the identification of discrete extra-SCN oscillators that tune circadian-modulated behaviors has been controversial and challenging^5^,^10,11^. For instance, cycling clock gene expression is observed broadly in neurons across the brain^5,12,13,14^, and it is unclear whether all, or a subset, of these cells function in promoting clock-dependent behavioral rhythms. In addition, local oscillators are thought to exhibit cycling electrical activity^14,15^, but there is a dearth of genetic markers that define electrically rhythmic neuronal clusters.

At the molecular level, it is well-established that the timekeeping mechanism comprises a transcriptional-translational feedback loop of core clock genes (e.g., *Clock*, *Bmal1*, *Period1/2*, *Cryptochrome*)^16,17^. However, the molecular mechanisms acting downstream of the clock to generate rhythmic outputs remain poorly understood. We previously identified the clock output molecule WIDE AWAKE (WAKE) in *Drosophila* and characterized its role in clock neurons, where it upregulates ion channels/pumps at night to promote rhythmic excitability^18,19^. A single homolog of WAKE is present in mice (mWAKE/ANKFN1), which is expressed in the SCN and a distributed set of brain regions, many of which exhibit cycling clock gene expression^5,14,20,21^. Because daily rhythms of electrical activity and clock gene expression are thought to be requisite features of extra-SCN oscillators^5,14,15^, we hypothesized that mWAKE defines neural circuits that function as circadian oscillators in the mouse brain.

Here, we show that mWAKE-expressing neurons are enriched in a subregion of the lateral amygdala (LA) and that impairing CLOCK function or excitatory signaling in these cells eliminates Period2 (Per2) cycling throughout the LA. At the cellular level, we find that mWAKE-positive, but not mWAKE-negative, LA neurons exhibit rhythmic excitability and that mWAKE promotes this electrical cycling by upregulating a Ca^2+^-dependent K^+^ (K_Ca_) current specifically at night. Rather than modulate a single behavior, our findings reveal that mWAKE-positive LA neurons coordinate sensory perception and emotional state in a time-dependent manner, thus regulating a circadian-dependent behavioral “module.” Moreover, we demonstrate that CLOCK function and mWAKE itself are required in mWAKE-positive LA neurons to generate these behavioral rhythms. Taken together, these data reveal the regional and cellular identity of an extra-SCN brain oscillator that regulates multiple clock-dependent behaviors and organizes cellular rhythms across brain regions.

## RESULTS

### mWAKE labels an LA subregion with rhythmic clock gene expression

We hypothesized that mWAKE labels brain subregions that represent brain oscillators and focused on the lateral nucleus of the amygdala (LA), because of its importance in emotional processing^22,23^ and the relatively strong mWAKE expression in this region^20^. Neuronal tracing experiments and lesion studies suggest the presence of distinct dorsal and ventral LA subregions^24–27^. This categorization was recently refined by a single cell RNA-Seq (scRNA-seq) analysis of the basolateral amygdala (BLA), which defined two major cell groups within the LA: a cluster labeled by *Myl4* in the anterior/dorsal subregion (adLA) and a cluster marked by *Otof* in the posterior/ventral subregion (pvLA)^28^. To characterize mWAKE expression in these LA subregions, we used two transgenic mouse lines^20,29^. In *mWake^Cre^* mice, exon 5 of *mWake* is replaced by a tdTomato-P2A-Cre cassette, which simultaneously labels *mWake*-positive cells and generates a loss-of-function allele (Figure 1A). In the second transgenic line, a V5 tag is fused to the C-terminus of mWAKE (*mWake^V^*^5^) (Figure 1B). Examination of native tdTomato signal in *mWake*^(Cre/+)^ mice and anti-V5 signal in *mWake*^(V5/V5)^ mice revealed that mWAKE^+^ cells are present in both the adLA and pvLA, but are markedly enriched in the former region (strong tdTomato signal in pvLA reflects expression in processes) (Figure 1C). Additional mWAKE expression is found in the central amygdala (CeA) and the pericapsular region outside the LA (Figure 1C).

**Figure 1.**
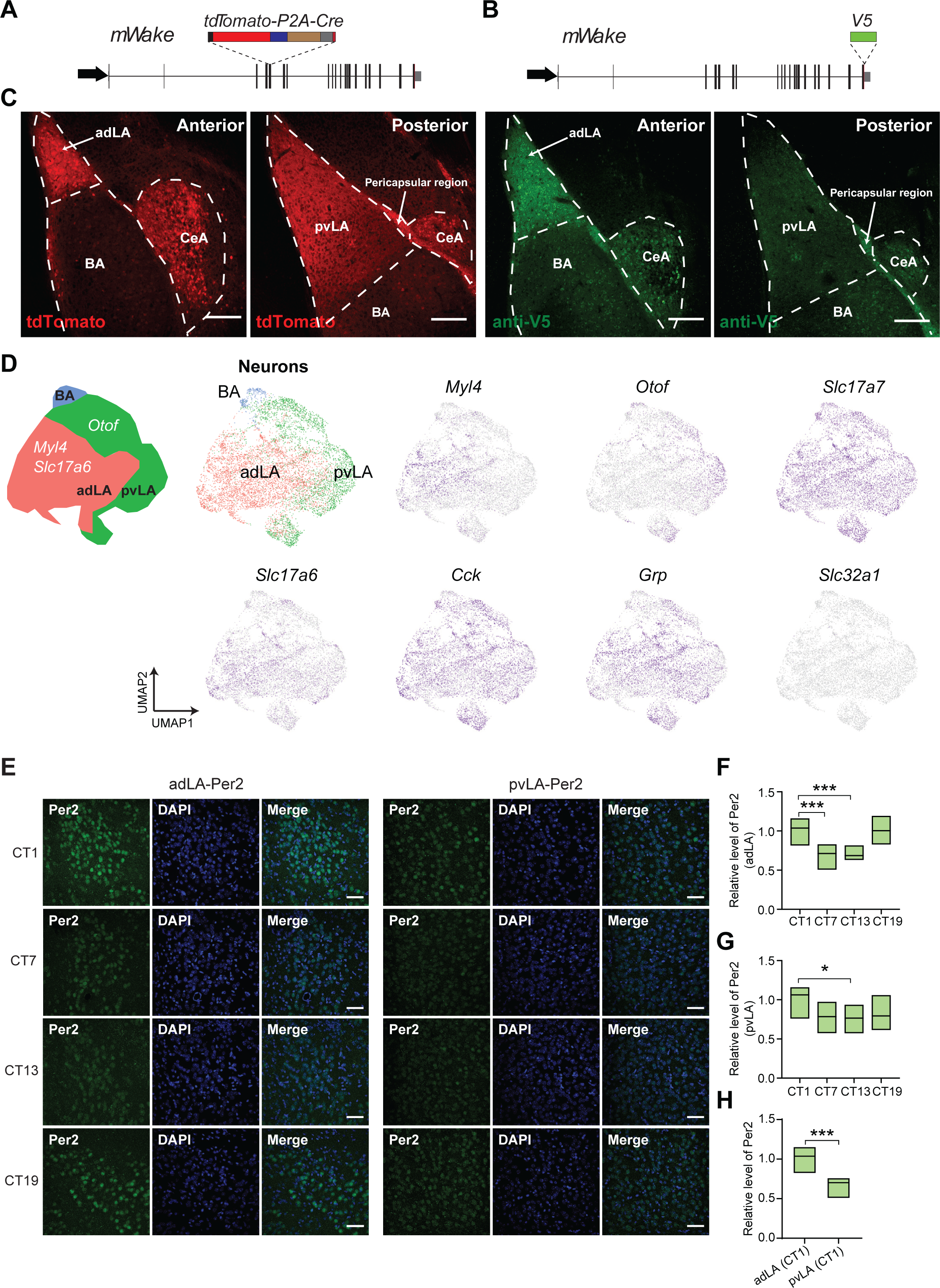
LA^mWake^ neuron identity and core clock rhythms. **(A)** Schematic for the *mWake^(Cre)^* transgene showing the genomic locus of *mWake* and replacement of exon 5 with a *tdTomato-P2A-Cre-stop* cassette. **(B)** Schematic for the *mWake^(V^*^5^*^)^* transgene showing the genomic locus of *mWake* and insertion of an in-frame V5 tag at the C-terminus. **(C)** Representative confocal images of anterior and posterior LA showing native tdTomato signal in an *mWake^(Cre/+)^*mouse (left) and anti-V5 immunostaining in an *mWake^(V^*^5^*^/V^*^5^*^)^* mouse (right), with “BA” (basal amygdala), “CeA” (central amygdala), “adLA” (anterior dorsal LA), “pvLA” (posterior ventral), and “Pericapsular” regions indicated. Scale bars represent 200 μm. **(D)** Schematic (left) and UMAP (Uniform Manifold Approximation and Projection) plots (right) showing 2 primary *mWake-*positive neuronal clusters in the LA (adLA and pvLA) and distribution of various marker genes across these clusters. Abbreviations: *Cck* (*cholecystokinin*), *GRP* (*gastrin-releasing peptide*), *Myl4* (*myosin light chain 4*), *Otof* (*otoferlin*), *Slc17a7* (*VGLUT1*), *Slc17a6* (*VGLUT2*), and *Slc32a1* (*VGAT*). **(E)** Representative confocal images of Per2 immunostaining in the adLA and pvLA regions at CT1, CT7, CT13 and CT19 in wild-type (WT) mice. DAPI and merged channels are also shown. Scale bars denote 50 μm. **(F** and **G)** Relative levels of Per2 intensity in the adLA region (**F**) (represented as a fold-change relative to CT1) at CT1 (n=23), CT7 (n=21), CT13 (n=22), and CT19 (n=18) and pvLA region **(G)** at CT1 (n=17), CT7 (n=13), CT13 (n=17), and CT19 (n=13) in WT mice. Data represented as simplified boxplots showing median with 1^st^ and 3^rd^ quartile boxes. n represents total number of sections, collected from 3-4 mice at each time point; Kruskall-Wallis test with post-hoc Dunn’s. **(H)** Relative levels of Per2 intensity in adLA and pvLA regions at CT1 condition (represented as a fold-change relative to adLA). Data represented as simplified boxplots showing median with 1^st^ and 3^rd^ quartile boxes; Kruskall-Wallis test with post-hoc Dunn’s. Data in (**H**) are from the same dataset shown in (**F** and **G**). In this figure and the following, error bars represent SEM and “*”, “**”, and “***” denote *P*<0.05, *P*<0.01, and *P*<0.001, respectively.

To investigate the identity of LA^mWAKE^ neurons, we performed both scRNA-seq and single nucleus RNA-Seq (snRNA-seq) from LA tissue of *mWake*^(Cre/+)^ mice (Figure 1D and S1A). LA^mWAKE^ cells were primarily neuronal, were found in both adLA and pvLA subpopulations, and express *Slc17a7*/*VGLUT1*, *cholecystokinin* (*CCK*), *gastrin-releasing peptide* (*GRP*), and, to a lesser extent, *Slc17a6/VGLUT2* (Figure 1D). In contrast, *Slc32A1/VGAT* was not expressed in these neurons (Figure 1D). Of note, we did not detect the presence of a distinctive neuronal marker expressed exclusively within mWAKE-positive neurons located in the LA, other than mWAKE itself. However, we did observe slightly elevated expression levels of K^+^ channel genes, including *Kcnh5* and *Kcnh1*, as well as Ca^2+^ channel genes, including *Cacna1a* and *Cacna1g*, in mWAKE-positive neurons when compared to mWAKE-negative neurons (Tables S1 and S2). These distinctions were more pronounced within the adLA subpopulation, where most mWAKE-positive neurons are located, compared to the pvLA populations and the pericapsular area. RNAscope *in situ* hybridization experiments in adLA confirmed *mWake* expression in this subregion in wild-type (WT) mice and co-labeling of *VGLUT2* and *GRP* (but not *VGAT*) with *tdTomato* in *mWake*^(Cre/+)^ mice (Figure S1B and S1C). To further verify the excitatory nature of LA^mWAKE^ neurons in adLA, we performed immunostaining using anti-CaMKIIα and anti-GAD67 antibodies and found that ∼80% of these cells express CaMKIIα, while none appeared to express GAD67 (Figure S1D and S1E).

Previous work has shown that levels of the core clock molecule Per2 cycle in the BLA^5,30,31^. To address whether Per2 cycling is observed in the LA, and specifically the adLA and pvLA subregions, we performed anti-Per2 immunostaining in WT mice at different times under constant darkness (DD). Our data suggest that, in the adLA, Per2 levels are higher at CT19 (Circadian Time) and CT1 (Figure 1E and 1F), roughly matching the phase of Per2 cycling previously shown for the BLA^30,31^. Although Per2 also cycles in the pvLA (Figure 1E and 1G), the overall level of Per2 expression in the pvLA is significantly lower than in the adLA at CT1 (Figure 1E and 1H). Together, these findings show that mWAKE labels a subset of excitatory neurons in the LA and is enriched in a subregion that demonstrates rhythmic clock gene expression.

### adLA^mWAKE^ neurons organize regional cellular rhythms

We next asked whether adLA^mWAKE^ neurons regulate clock gene rhythms in the LA. To address this question, we sought to selectively disrupt core clock function in adLA^mWAKE^ neurons. Thus, we generated a CLOCK-dominant negative (Clock-DN) construct based on a validated design used in *Drosophila*, in which the DNA-binding basic domain was deleted while leaving the protein-interaction helix-loop-helix domain largely intact (Figure 2A)^32^. Using this construct, we produced a Cre-dependent Clock-DN virus (AAV-DIO-Clock-DN-P2A-EYFP) (Figure 2A). To validate the ability of this virus to impair CLOCK function, we injected it into the SCN of *VGAT*^(Cre/Cre)^ mice (Table S3) and assayed locomotor activity rhythms using a wheel-running assay (Figure S2A and S2B). Expression of AAV-DIO-Clock-DN-P2A-EYFP virus in GABAergic SCN neurons significantly reduced rhythm strength, without significantly altering period length, compared to control AAV-DIO-EYFP viral injections (Figure S2C-S2G).

**Figure 2.**
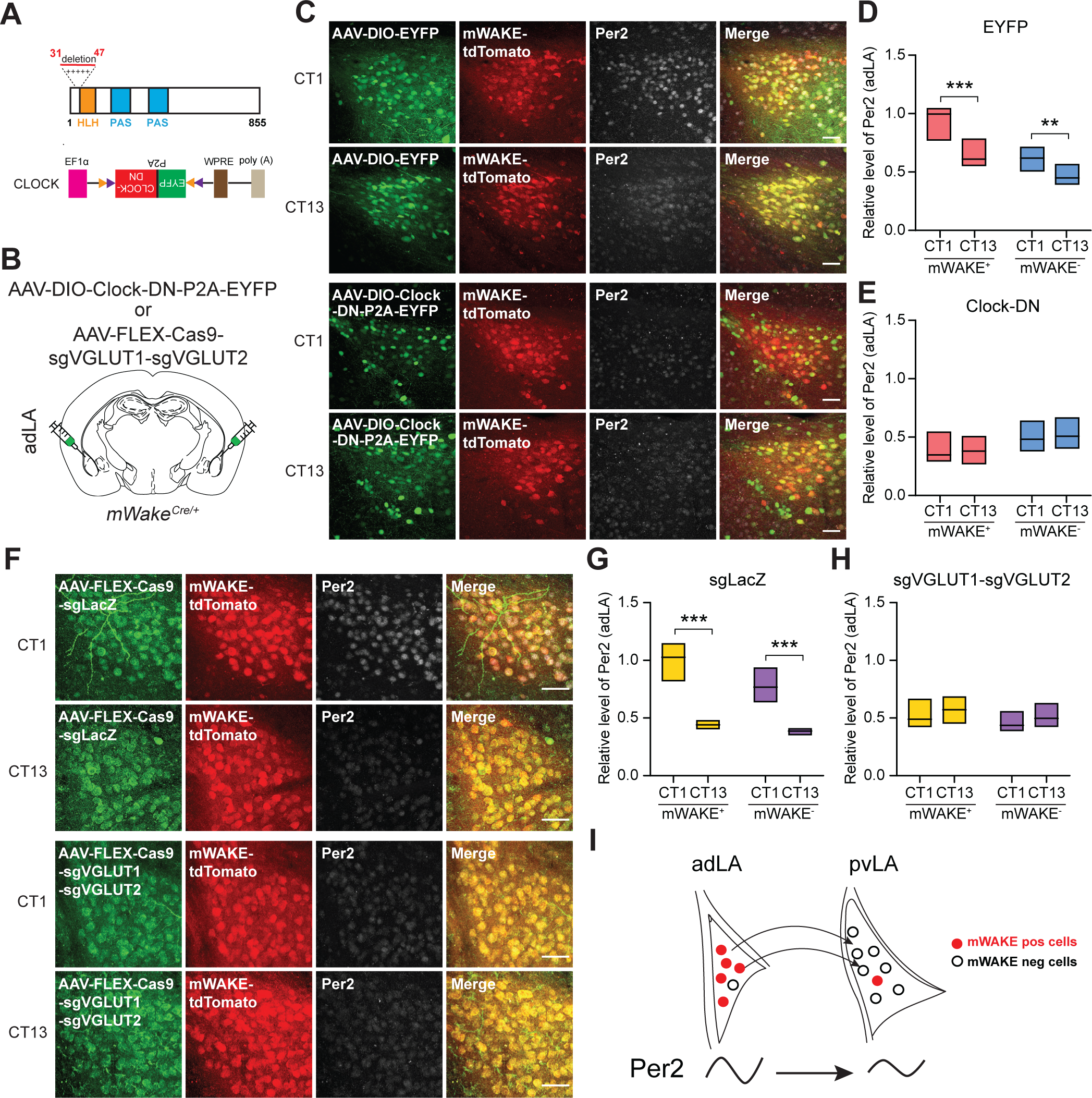
adLA^mWAKE^ neurons organize molecular rhythms in the LA. **(A)** Schematics illustrating design of the Clock dominant negative (Clock-DN) construct (top) and Cre-dependent Clock-DN virus (bottom). (top) 17 aa from the DNA-binding basic region of Clock (Lys31-Arg47) are deleted in the Clock-DN construct. The helix-loop-helix (HLH) and PAS domains are indicated by orange and blue boxes, respectively. (bottom) The AAV-DIO-Clock-DN-P2A-EYFP sequence includes an EF1α promoter, an inverted Clock-DN-P2A-EYFP cassette flanked by a pair of loxP (orange) and lox2722 (purple) sites with inward orientation, a woodchuck hepatitis virus post-transcriptional regulatory element (WPRE), and polyA signal. **(B)** Schematic image showing the injection of AAV-DIO-Clock-DN-P2A-EYFP or AAV-FLEX-Cas9-sgVGLUT1-sgVGLUT2 virus into the adLA of *mWake^(Cre/+)^* mice. **(C)** Representative confocal images of native EYFP and tdTomato or Per2 immunostaining in the adLA region at CT1 vs CT13 in *mWake^(Cre/+)^* mice with either AAV-DIO-EYFP (top) or AAV-DIO-Clock-DN-P2A-EYFP (bottom) injected bilaterally into the adLA. Merged channels are also shown. Scale bars denote 50 μm. **(D** and **E)**, Relative levels of Per2 intensity in mWAKE-positive (red) and mWAKE-negative (cyan) cells in adLA at CT1 vs CT13 in *mWake^(Cre/+)^*mice injected with either AAV-DIO-EYFP **(D)** (CT1, n=21; CT13, n=13) or AAV-DIO-Clock-DN-P2A-EYFP (**E**) (CT1, n=21; CT13, n=12) injected bilaterally into the LA. Data represented as simplified boxplots showing median with 1^st^ and 3^rd^ quartile boxes. n represents total number of sections, collected from 3-4 mice for each condition. Data represented as a fold-change relative to the signal for mWAKE-positive cells under control CT1 condition; Mann-Whitney U tests with Bonferroni correction. **(F)** Representative confocal images of native tdTomato or Cas9 and Per2 immunostaining in the adLA region at CT1 vs CT13 in *mWake^(Cre/+)^* mice with either control AAV-FLEX-Cas9-sgLacZ or AAV-FLEX-Cas9-sgVGLUT1-sgVGLUT2 injected bilaterally into the adLA. Merged channels are also shown. Scale bars denote 50 μm. **(G** and **H)** Relative levels of Per2 intensity in mWAKE-positive (yellow) and mWAKE-negative (purple) cells in the adLA region at CT1 vs CT13 in *mWake^(Cre/+)^*mice with either AAV-FLEX-Cas9-sgLacZ (CT1, n=12; CT13, n=13) (**G**) or AAV-FLEX-Cas9-sgVGLUT1-sgVGLUT2 (CT1, n=28; CT13, n=26) (**H**) injected bilaterally into the adLA. Data represented as simplified boxplots showing median with 1^st^ and 3^rd^ quartile boxes. n represents number of sections, and 4 mice were used for each condition. Data represented as a fold-change relative to the signal in control mWAKE-positive neurons at CT1; Mann-Whitney U tests with Bonferroni correction. **(I)** Model. mWAKE-positive neurons are enriched in the adLA, which exhibits Per2 cycling. adLA neurons project to the pvLA, and this excitatory signaling is required for Per2 rhythms in the pvLA.

We next bilaterally injected AAV-DIO-Clock-DN-P2A-EYFP or AAV-DIO-EYFP into the adLA of *mWake*^(Cre/+)^ mice (Figure 2B and Table S3) and then performed immunostaining for Per2 in mWAKE-positive and mWAKE-negative cells in adLA and pvLA. In control mice, following injection of AAV-DIO-EYFP virus, Per2 levels cycled in both mWAKE-positive and mWAKE-negative neurons, with higher Per2 signal at CT1 compared to CT13 (Figure 2C and 2D). Strikingly, selective expression of Clock-DN in mWAKE-positive neurons in adLA not only abolished cycling of Per2 in a cell-autonomous manner, but also did so in both mWAKE-negative neurons in the adLA (Figure 2C and 2E) and the pvLA (Figure S3A-S3D). We next examined the effects of impairing CLOCK function in mWAKE-negative neurons in the pvLA region. To do this, we generated an AAV-DO-Clock-DN-EYFP virus to drive Clock-DN expression in a “Cre-off”-dependent manner and injected this virus bilaterally into the pvLA of *mWake*^(Cre/+)^ mice (Figure S3E and Table S3). Interestingly, Per2 cycling was maintained in mWAKE-positive and even in mWAKE-negative cells (where CLOCK function was directly impaired) in the pvLA (Figure S3F and S3G), suggesting that Per2 rhythms in the pvLA depend on extrinsic inputs.

Excitatory glutamate signaling has previously been shown to affect Per2 transcription^33,34^. Thus, we hypothesized that adLA^mWAKE^ neurons coordinate Per2 rhythms in the pvLA through glutamate signaling. To test this possibility, we performed CRISPR-mediated knockout of VGLUT1 and VGLUT2 in adLA^mWAKE^ cells by injecting AAV-FLEX-Cas9-sgVGLUT1-sgVGLUT2 virus^35^ into the adLA of *mWake*^(Cre/+)^ mice and performed Per2 immunostaining in both the adLA and pvLA (Figure 2B, S3H and Table S3). As predicted, Per2 cycling was lost in both mWAKE-positive and mWAKE-negative neurons in both adLA (Figure 2F and 2H) and pvLA (Figure S3I and S3K). In contrast, expressing control AAV-FLEX-Cas9-sgLacZ virus in adLA^mWAKE^ cells did not affect Per2 rhythms in either the adLA (Figure 2F and 2G) or pvLA (Figure S3I and S3J). Consistent with a role for the adLA region coordinating Per2 rhythms in the pvLA, we found substantial projections emanating from adLA^mWAKE^ neurons to the pvLA region (Figure S3L). In contrast, no clear projections were observed from pvLA^VGLUT1^ neurons to the adLA region (Figure S3M). Together, these findings suggest a hierarchical relationship between adLA and pvLA regions, whereby adLA^mWAKE^ neurons signal to the pvLA and organize local Per2 rhythms throughout the LA (Figure 2I).

### mWAKE expression cycles and defines a rhythmically excitable neuronal cluster in the LA

Based on our previous work on WAKE/mWAKE in *Drosophila* clock neurons^18,19^ and mouse dorsomedial hypothalamus (DMH)^29^, we next asked whether mWAKE acts to inhibit neuronal excitability at night in the LA. We first performed Western blots using LA tissue from *mWake^(V^*^5^*^/V^*^5^*^)^* mice at different time points in DD. These data demonstrated that mWAKE levels cycle in the LA under constant conditions, peaking at CT13 (Figure 3A and 3B). Next, we examined whether the electrical activity of adLA^mWAKE^ neurons also cycles. We performed whole-cell patch-clamp slice recordings of mWAKE-positive cells from the adLA region in *mWake^(Cre/+)^* vs mutant *mWake^(Cre/Cre)^* mice (Figure S4A). Because we did not observe spontaneous firing, we examined intrinsic electrical excitability by measuring spiking frequency in response to current injections. In control *mWake^(Cre/+)^* mice, the intrinsic excitability of adLA^mWAKE^ neurons was reduced at night (Zeitgeber Time 12-14/ZT12-14) compared to the day (ZT0-2). In contrast, this cycling of intrinsic excitability was lost in mutant adLA^mWAKE^ neurons (*mWake^(Cre/Cre)^*), due to an increase in excitability of these cells at night (Figure 3C and 3D).

**Figure 3.**
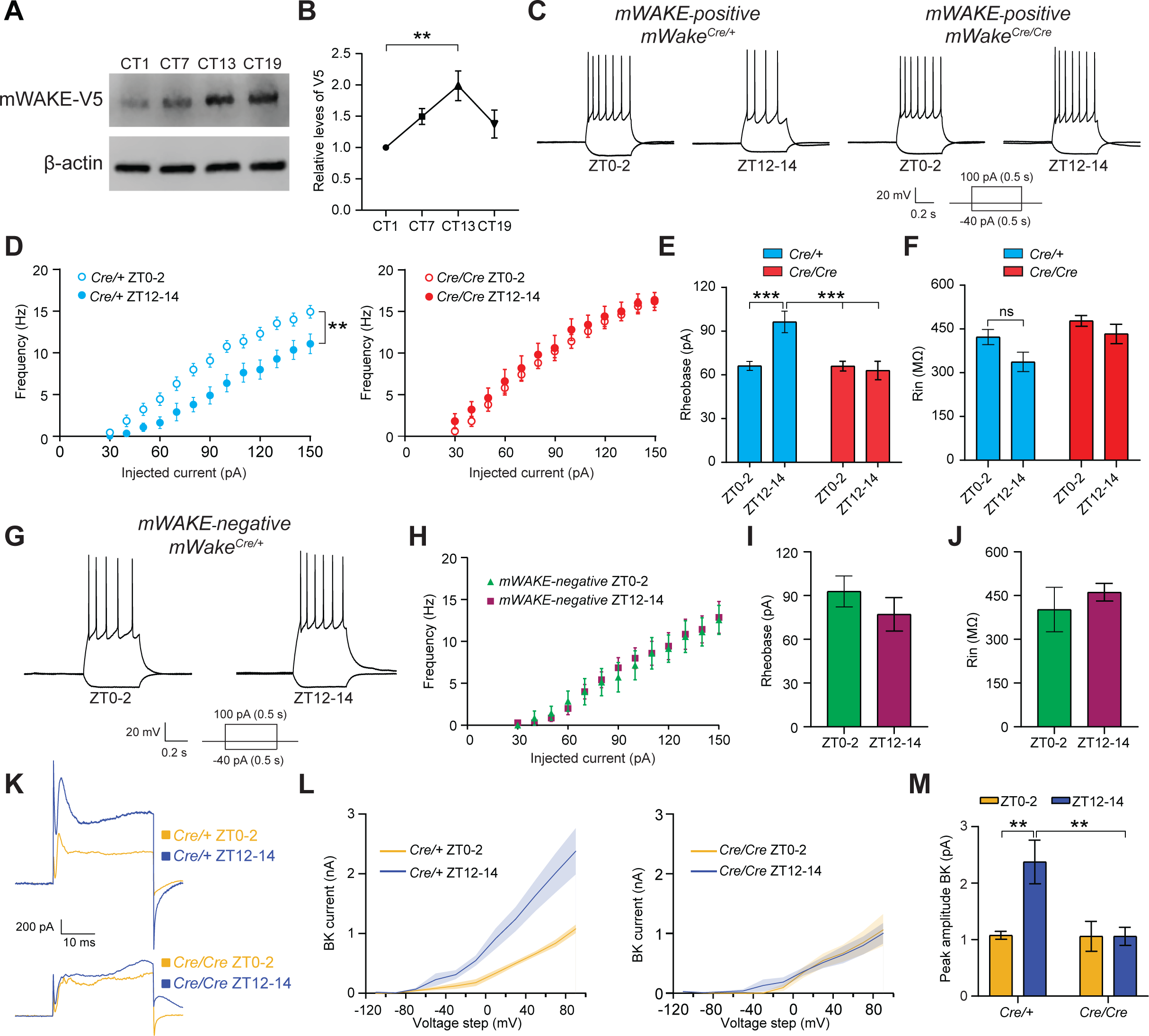
Cycling of mWAKE and neuronal excitability in adLA^mWAKE^ neurons. **(A and B)** Representative immunoblot (**A**) and relative levels of mWAKE-V5 normalized to a β-actin loading control (**B**) from Western blot analyses of *mWake^(V^*^5^*^/V^*^5^*^)^* LA tissue using anti-V5 antibodies at CT1, CT7, CT13, and CT19 (n=7 mice for all time points); one-way ANOVA with post-hoc Dunnett’s test. **(C)** Representative membrane potential traces from whole-cell patch-clamp recordings of adLA^mWAKE^ neurons following current injection of −40 pA and 100 pA in *mWake^(Cre/+)^* and *mWake^(Cre/Cre)^* mice at ZT0-2 and ZT12-14. **(D)** *f-I* curves for adLA^mWAKE^ neurons from *mWake^(Cre/+)^* (cyan, left) or *mWake^(Cre/Cre)^* (red, right) mice at ZT0-2 (open circles) vs ZT12-14 (closed circles) (n=10-13 cells for each group from 3-4 animals); two-way ANOVA with post-hoc Sidak. **(E** and **F)** Rheobase (minimum current amplitude to trigger an action potential) (**E**) and input resistance (Rin) (**F**) for adLA^mWAKE^ neurons in *mWake^(Cre/+)^* (cyan) or *mWake^(Cre/Cre)^* (red) mice at ZT0-2 vs ZT12-14. Data are from the same cells shown in (**D**); two-way ANOVA with post-hoc Sidak. **(G)** Representative membrane potential traces from whole-cell patch-clamp recordings of mWAKE-negative LA neurons following current injection of −40 pA or 100 pA at ZT0-2 and ZT12-14. **(H**-**J)** *f-I* curves (**H**), rheobase (**I**), and input resistance (**J**) for mWAKE-negative adLA neurons from *mWake^(Cre/+)^* mice at ZT0-2 (green) vs ZT12-14 (magenta) (n=7 cells for each group from 3 animals); two-way ANOVA post-hoc Sidak. **(K)** Representative traces of BK current in adLA^mWAKE^ neurons from *mWake^(Cre/+)^* (top) and *mWake^(Cre/Cre)^* (bottom) at ZT0-2 (orange) vs ZT12-14 (blue). **(L)** Average BK current at different voltage steps in adLA^mWAKE^ neurons from *mWake^(Cre/+)^*(left) and *mWake^(Cre/Cre)^* (right) at ZT0-2 (orange) vs ZT12-14 (blue). (n=7-10 cells from 3 animals). Shading denotes SEM. **(M)** Peak amplitude of BK current in adLA^mWAKE^ neurons from *mWake^(Cre/+)^* and *mWake^(Cre/Cre)^* at ZT0-2 (orange) vs ZT12-14 (blue). Data are from the same dataset as in (**L**); two-way ANOVA post-hoc Sidak.

In line with these findings, rheobase, the amount of current needed to induce the initial spike, was increased at night compared to the daytime in control adLA^mWAKE^ neurons, and this cycling of rheobase was absent in mutant adLA^mWAKE^ neurons (Figure 3E). In contrast, we did not observe significant differences in day/night cycling in control vs mutant adLA^mWAKE^ neurons for measures of passive membrane properties, such as input resistance (Rin) and resting membrane potential (RMP) (Figure 3F and S4B). We repeated these experiments for mWAKE-negative neurons in the adLA region (Figure S4A). Interestingly, these data revealed a lack of cycling of intrinsic excitability for these cells (Figure 3G and 3H). In addition, rheobase, Rin, and RMP were similar between daytime and nighttime (Figure 3I, 3J and S4C). Taken together, these data suggest that mWAKE levels cycle in the LA to suppress excitability at night and that mWAKE defines adLA neurons that are intrinsically rhythmic.

How the circadian clock promotes rhythmic excitability outside the SCN is poorly understood^15^. Our previous work in *Drosophila* suggested that WAKE acts to upregulate specific ion channels and receptors at night^18,19^. One of these ion channels in *Drosophila* is Slowpoke, a Ca^2+^-activated voltage-gated K^+^ channel, whose mammalian homolog is BK/KCa1.1^19^. We thus hypothesized that mWAKE upregulates BK current at night in the adLA. We performed whole-cell voltage-clamp slice recordings of adLA^mWAKE^ neurons in *mWake^(Cre/+)^*or *mWake^(Cre/Cre)^* mice and recorded BK current at ZT0-2 vs ZT12-14. We found that BK current amplitude cycled in control adLA^mWAKE^ neurons, with greater BK current at night vs the day. This cycling was lost in mutant adLA^mWAKE^ neurons, with BK current reduced both during the day and night (Figure 3K-3M). These data suggest that mWAKE promotes BK channel activity at night to inhibit neuronal excitability in a rhythmic manner.

### adLA^mWAKE^ neurons project broadly outside the amygdala

Because our data suggested that adLA^mWAKE^ neurons define an extra-SCN oscillator, we next investigated whether they generate rhythmic behaviors. Because a clear behavioral role has not been assigned to the pvLA region, we sought to identify potential behaviors that adLA^mWAKE^ neurons regulate, by first assessing the projection pattern of adLA^mWAKE^ neurons. We injected AAV-FLEX-EGFP into the adLA of *mWake^(Cre/+)^* mice unilaterally and examined native fluorescence by confocal microscopy (Table S3). In addition to dense projections to the basal amygdala (BA), our experiments revealed prominent projections to secondary somatosensory cortex (S2), the nucleus accumbens (NAc) core, the bed nucleus of the stria terminalis (BNST), auditory cortex (ACx), and substantia nigra (SN) (Figure 4A). To demonstrate that these projections represent synaptic terminals, we repeated the experiments using AAV-DIO-ChR2-EYFP (Table S3) and confirmed EYFP expression in S2, NAc core, BNST, ACx, and SN regions. In addition, we identified projections to many regions throughout the cortex, subcortical regions, and brainstem (Figure S5A and S5B). Taken together, these data suggest that adLA^mWAKE^ neurons send projections to a wide array of extra-amygdala regions, raising the possibility that they promote a variety of rhythmic behaviors.

**Figure 4.**
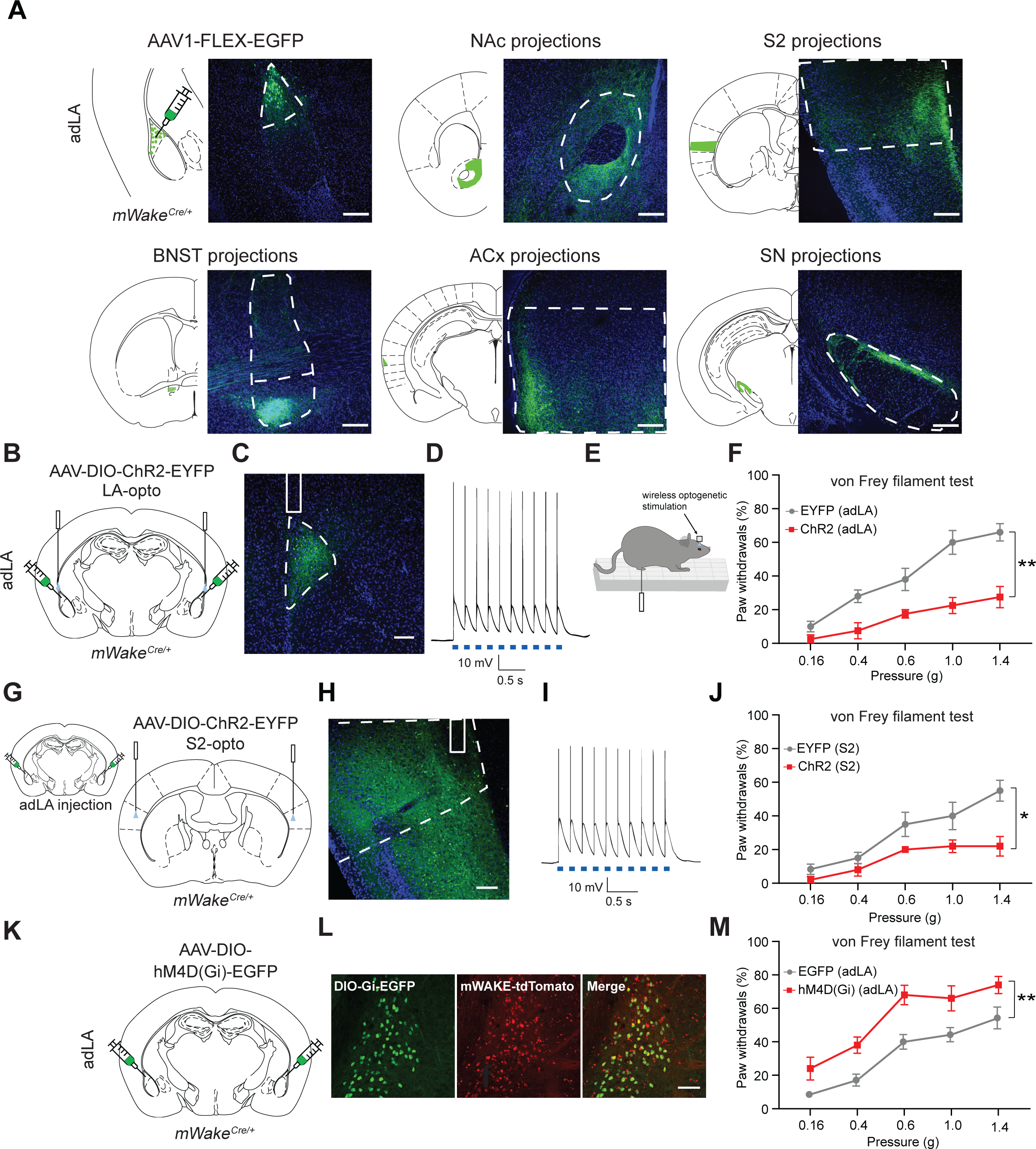
adLA^mWAKE^ neurons inhibit mechanical sensitivity via projections to S2. **(A)** Schematics and representative confocal images of native GFP signal following injection of AAV1-FLEX-EGFP virus into the adLA of an *mWake^(Cre/+)^*mouse showing cell bodies in adLA and terminal projections in NAc core (nucleus accumbens core), S2 (secondary somatosensory cortex), BNST (bed nucleus of the stria terminals), ACx (auditory cortex), and SN (substantia nigra). Scale bars represent 200 μm. **(B)** Schematic showing bilateral injections of AAV-DIO-ChR2-EYFP virus and optical fiber implantation into the adLA of *mWake^(Cre/+)^* mice. **(C)** Representative confocal image of YFP native fluorescence in the adLA (dashed outline) following injection of the virus described in (**B**). Location of optical fiber (solid line) is shown. Scale bar represents 100 μm. **(D)** Representative membrane potential trace from a whole-cell patch-clamp recording of an adLA^mWAKE^ neuron showing action potentials triggered by adLA optogenetic stimulation. Blue boxes indicate 5 ms blue light pulses. **(E)** Schematic depicting the mechanical sensitivity assay applying Von Frey filaments to deliver touch stimuli of different strengths, while activating the circuit using wireless optogenetic stimulation. **(F)** Average hind foot withdrawal (%) in response to mechanical stimuli delivered using von Frey filaments of different strengths in the presence of 20 Hz optogenetic stimulation of adLA^mWAKE^ neurons in *mWake^(Cre/+)^* mice injected with AAV-DIO-ChR2-EYFP (n=4, red) or control AAV-DIO-EYFP (n=5, gray); two-way ANOVA with post-hoc Sidak. **(G)** Schematics showing bilateral injections of AAV-DIO-ChR2-EYFP into the adLA (left) and optical fiber implantation into the S2 region (right) of *mWake^(Cre/+)^* mice. **(H)** Representative confocal image of YFP native fluorescence in S2 (dashed outline) for the mouse shown in (**G**). Location of optical fiber (solid line) is shown. Scale bar represents 100 μm. **(I)** Representative membrane potential trace from a whole-cell patch-clamp recording of an S2 neuron near adLA^mWAKE^ terminals showing action potentials triggered by S2 optogenetic stimulation. Blue boxes indicate 5 ms blue light pulses. **(J)** Average hind foot withdrawal (%) in response to mechanical stimuli delivered using von Frey filaments of different strengths in the presence of 20 Hz optogenetic stimulation of the S2 terminals of adLA^mWAKE^ neurons in *mWake^(Cre/+)^* mice injected with AAV-DIO-ChR2-EYFP (n=5, red) or control AAV-DIO-EYFP (n=6, gray); two-way ANOVA with post-hoc Sidak. **(K)** Schematic showing bilateral injections of AAV-DIO-hM4D(Gi)-EGFP virus in the adLA of *mWake^(Cre/+)^* mice. **(L)** Representative confocal images of the LA showing co-labeling of virally-expressed GFP fluorescence and native tdTomato fluorescence for the animal shown in (**K**). Scale bar represents 100 μm. **(M)** Average hind foot withdrawal (%) in response to mechanical stimuli delivered using von Frey filaments of different strengths following 10 mg/kg CNO injection in *mWake^(Cre/+)^*mice with AAV-DIO-hM4D(Gi)-EGFP (n=5, red) or AAV-DIO-EYFP (n=7, gray) injected into the adLA; two-way ANOVA with post-hoc Sidak.

### adLA^mWAKE^ neurons inhibit mechanical sensitivity and promote anxiety

To address the functional role of adLA^mWAKE^ neurons, we performed optogenetic and chemogenetic manipulations and assessed different behaviors. Given the prominent adLA^mWAKE^ projections to S2 (Figure 4A, S5A, and S5B), we first asked whether these cells regulate touch and/or pain perception. We injected AAV-DIO-ChR2-EYFP virus or a control virus bilaterally into the adLA of *mWake^(Cre/+)^* mice (Table S3) and then performed optogenetic stimulation of the LA, while simultaneously applying von Frey filaments of differing strengths to the hind paw of the mouse in a blinded fashion. These experiments demonstrated that optogenetic activation of adLA^mWAKE^ neurons reduced the frequency of paw withdrawal to a range of stimuli, including both innocuous and painful (Figure 4B-4F). To confirm that this effect was mediated by adLA^mWAKE^➔S2 projections, we repeated these experiments but instead implanted the optical fiber in the S2 region (Figure 4G and 4H). Optogenetic stimulation of adLA^mWAKE^➔S2 terminals induced firing of downstream S2 neurons (Figure 4I), and the optogenetically-triggered excitatory post-synaptic currents (oEPSCs) in these downstream neurons were suppressed using ionotropic glutamate receptor blockers (NBQX and D-AP5) (Figure S6A). As expected, mechanical sensitivity was similarly reduced with optogenetic stimulation of adLA^mWAKE^➔S2 terminals (Figure 4J). To investigate the effects of inhibiting adLA^mWAKE^ neurons on cutaneous mechanical sensitivity, we used a chemogenetic approach by injecting AAV-DIO-hM4D(Gi)-EGFP or a control virus bilaterally into the adLA of *mWake^(Cre/+)^* mice (Table S3) and then administering CNO (Figure 4K and 4L). Chemogenetic inhibition of adLA^mWAKE^ neurons led to an increase in paw withdrawal frequency following mechanical stimulation (Figure 4M). These data indicate that adLA^mWAKE^ neurons inhibit mechanical sensitivity and demonstrate a role for LA neurons in directly tuning sensory perception. Because adLA^mWAKE^ neurons are excitatory, they likely signal to inhibitory interneurons in S2 to reduce perception of touch and pain.

adLA^mWAKE^ neurons also send dense projections to the NAc core. The NAc is well-known to function in motivation and reward^36–38^, but has also been implicated in regulating anxiety and aversive-related behaviors^37,39–42^. To examine whether adLA^mWAKE^➔NAc signaling is rewarding or aversive, we first performed place preference/avoidance assays. We expressed AAV-DIO-ChR2-EYFP virus or control virus in adLA^mWAKE^ neurons and found that mice spent less time on the side associated with optogenetic activation (Figure 5A-5C), suggesting that adLA^mWAKE^ neurons participate in aversive, not reward-associated, behaviors. We then injected AAV-DIO-ChR2-EYFP or control virus bilaterally into the LA of *mWake^(Cre/+)^* mice (Table S3) and performed photostimulation at terminals in the NAc core, which induced spiking in NAc core neurons (Figure 5D-5F). As expected, oEPSCs in these downstream NAc core neurons were inhibited by blockade of excitatory input (Figure S6B). Similarly, optogenetic activation of adLA^mWAKE^ terminals in the NAc core also induced place avoidance (Figure 5G), indicating that adLA^mWAKE^➔NAc signaling is aversive. We next asked whether the adLA^mWAKE^➔NAc core circuit promotes anxiety-like behaviors. To do this, we performed open-field (OF) and elevated plus-maze (EPM) tests. Optogenetic stimulation of adLA^mWAKE^➔NAc core terminals decreased time spent in the central zone in the OF test, as well as in the open arm in the EPM test, compared to animals with control virus injection (Figure 5H-5K). No differences in the OF and EPM tests were observed prior to and after optogenetic stimulation (Figure S6C and S6D). We also examined whether chemogenetic inhibition of adLA^mWAKE^ neurons affected anxiety-like behaviors but found that this manipulation did not affect time spent in the center zone in the OF test or in the open arm in the EPM test (Figure S6E and S6F), compared to mice injected with control virus. Taken together, these data suggest that adLA^mWAKE^ neurons inhibit touch sensitivity and promote anxiety, thus coordinating sensory perception and emotional state.

**Figure 5.**
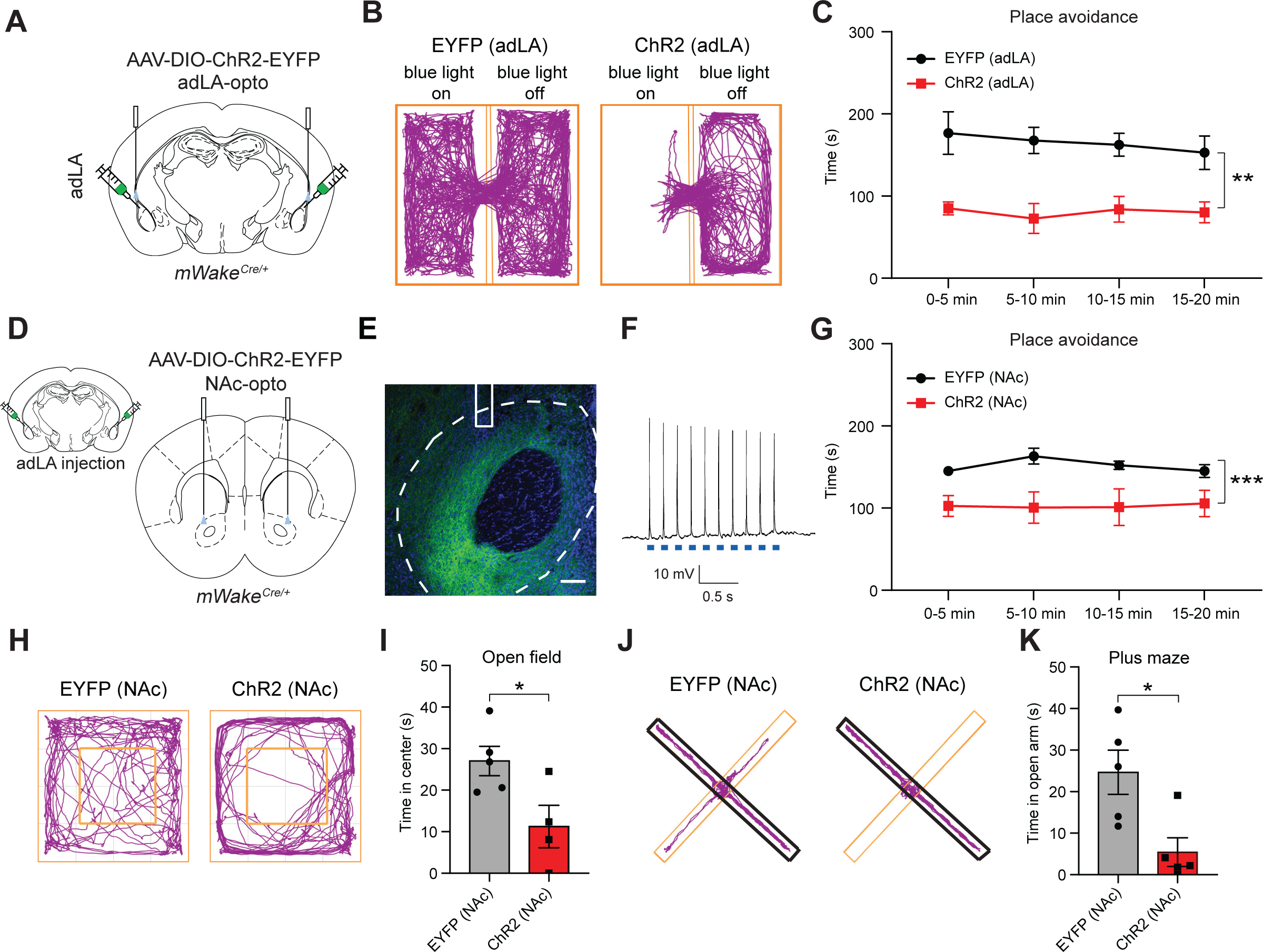
adLA^mWAKE^ neurons enhance anxiety via projections to NAc core. **(A)** Schematic showing bilateral injections of AAV-DIO-ChR2-EYFP virus and optical fiber implantation into the adLA of an *mWake^(Cre/+)^* mouse. **(B** and **C)** Representative locomotor activity tracks (**B**) and average time (s) spent in the LED light-stimulating side in 5 min bins (**C**) during a 20 min place preference/avoidance test during for the mice described in (**A**), injected with AAV-DIO-ChR2-EYFP (n=5, red) or control AAV-DIO-EYFP (n=6, black). 20 Hz optogenetic stimulation of adLA^mWAKE^ neurons was delivered when the mouse entered the “on” side and stopped when it entered the “off” side. **(D)** Schematics showing bilateral injections of AAV-DIO-ChR2-EYFP into the adLA (left) and optical fiber implantation into the NAc core region (right) of *mWake^(Cre/+)^* mice. **(E)** Representative confocal image of YFP native fluorescence in NAc core (dashed outline) for the mouse shown in (**D**). Location of optical fiber (solid line) is shown. Scale bar represents 100 μm. **(F)** Representative membrane potential trace from a whole-cell patch-clamp recording of a NAc core neuron near adLA^mWAKE^ terminals showing action potentials triggered by NAc core optogenetic stimulation. Blue boxes indicate 5 ms blue light pulses. **(G)** Average time (s) spent in the LED light stimulating side in 5 min bins during a 20 min place preference/avoidance test for *mWake^(Cre/+)^*mice injected with AAV-DIO-ChR2-EYFP (n=4, red) or control AAV-DIO-EYFP (n=4, black). 20 Hz optogenetic stimulation of the NAc terminals of adLA^mWAKE^ neurons was delivered when the mouse entered the “on” side and stopped when it entered the “off” side. **(H** and **I)** Representative locomotor activity tracks (**H**) and average time (s) spent in center zone (**I**) in open field assay during 5 min 20 Hz optogenetic stimulation of *mWake^(Cre/+)^* mice injected with AAV-DIO-ChR2-EYFP (n=4, red) or control AAV-DIO-EYFP (n=5, gray) into the adLA with optical fibers implanted in the NAc core. Center zone is indicated by yellow square; unpaired t-test, two-tailed. **(J** and **K)** Representative locomotor activity tracks (**J**) and average time (s) spent in open arm (**K**) during elevated plus maze assays during 5 min 20 Hz optogenetic stimulation of *mWake^(Cre/+)^* mice injected with AAV-DIO-ChR2-EYFP (n=5, red) or control AAV-DIO-EYFP (n=5, gray) into the adLA with optical fibers implanted in NAc. Closed and open arms indicated with thick black lines and thin yellow lines, respectively; unpaired t-test, two-tailed.

### CLOCK function is required in adLA^mWAKE^ neurons to generate rhythmic behaviors

We next used the AAV-DIO-Clock-DN-P2A-EYFP virus to investigate whether local clock function in the adLA^mWAKE^ neurons is required to regulate rhythms of mechanical sensitivity and anxiety (Figure 6A and Table S3). Previous work has generally suggested that pain sensitivity and anxiety-like behaviors cycle in a daily manner, but these studies have yielded mixed results and were usually performed under L:D conditions^43–47^. Thus, we asked whether, under constant conditions, WT mice exhibit cycling of mechanical sensitivity and anxiety-like behaviors. WT mice were more sensitive to mechanical stimuli during CT12-14 (their active period), compared to CT0-2 (Figure S7A). In addition, WT mice spent more time in the center zone in the OF test and in the open arm in the EPM test during CT12-14 (Figure S7B and S7C), suggesting that WT mice were less anxious during their active period at night. Similar cycling of cutaneous mechanical sensitivity and anxiety-like behaviors were also observed in control *mWake^(Cre/+)^* mice bilaterally injected with AAV-DIO-EYFP virus into the LA (Figure 6B, 6D, 6F, 6H, and 6J). In contrast, bilateral injection of the AAV-DIO-Clock-DN-P2A-EYFP virus in the adLA of *mWake^(Cre/+)^* mice resulted in the loss of daily rhythms of mechanical sensitivity (Figure 6C) and anxiety-like behavior (Figure 6E, 6G, 6I, and 6K), such that their behavioral responses during the subjective night resembled those seen during the subjective day. Together, these data argue that local clock function is required in LA^mWAKE^ neurons for cycling of sensory perception and emotional state and support the notion that these cells constitute an extra-SCN circadian oscillator.

**Figure 6.**
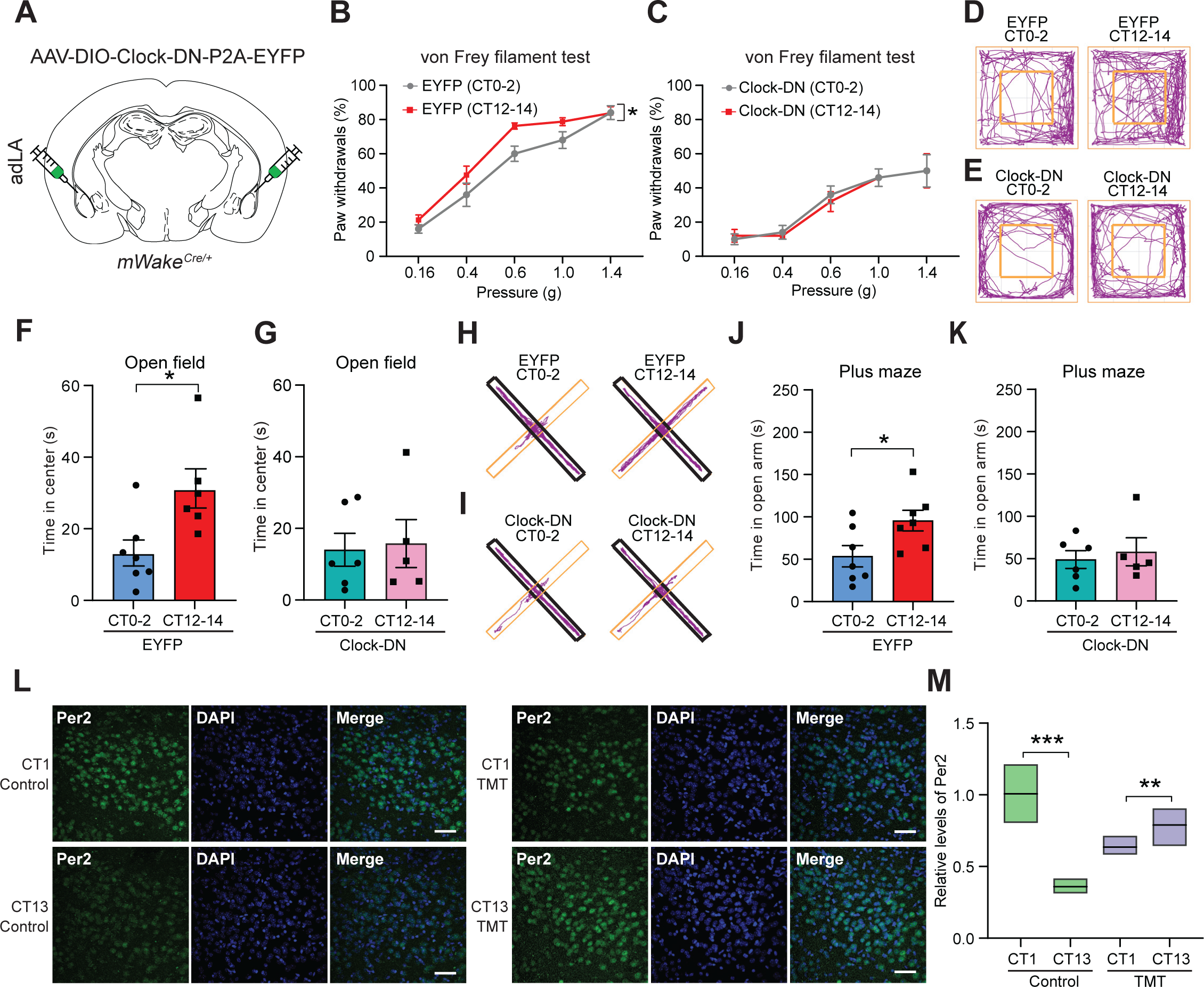
A local clock in adLA^mWAKE^ neurons mediates rhythmic regulation of mechanical sensitivity and anxiety. **(A)** Schematic showing bilateral injection of Cre-dependent Clock-DN virus (AAV-DIO-Clock-DN-P2A-EYFP) in the adLA of *mWake^(Cre/+)^* mice. **(B** and **C)** Average hind foot withdrawal (%) in response to mechanical stimuli delivered using von Frey filaments of different strengths at CT0-2 (gray) or CT12-14 (red) for *mWake^(Cre/+)^* mice injected with control AAV-DIO-EYFP (**B**) (CT0-2, n=5; CT12-14, n=8) or AAV-DIO-Clock-DN-P2A-EYFP (**C**) (CT0-2, n=5; CT12-14, n=5) into the adLA; two-way ANOVA with post-hoc Sidak. **(D** and **E)** Representative locomotor activity tracks in open field assays at CT0-2 or CT12-14 for *mWake^(Cre/+)^* mice injected with control AAV-DIO-EYP (EYFP) (**D**) or AAV-DIO-Clock-DN-P2A-EYFP (Clock-DN) (**E**) into the adLA. Center zone is indicated by yellow square. **(F** and **G)** Average time spent in center zone in open field assay at CT0-2 vs CT12-14 for *mWake^(Cre/+)^* mice injected with control AAV-DIO-EYFP (**F**) (EYFP: CT0-2, n=7 (cyan); CT12-14, n=6 (red)) or AAV-DIO-Clock-DN-P2A-EYFP (**G**) (Clock-DN: CT0-2, n=6 (cyan-green); CT12-14, n=5 (pink)) into the adLA; unpaired t-test, two-tailed. **(H** and **I)** Representative locomotor activity tracks for elevated plus maze assays at CT0-2 vs CT12-14 for *mWake^(Cre/+)^* mice injected with control AAV-DIO-EYFP (**H**) (EYFP) or AAV-DIO-Clock-DN-P2A-EYFP (**I**) (Clock-DN) into the adLA. Closed and open arms indicated with thick black lines and thin yellow lines, respectively. **(J** and **K)** Average time (s) spent in open arm during elevated plus maze assays at CT0-2 vs CT12-14 for *mWake^(Cre/+)^* mice injected with control AAV-DIO-EYFP (**J**) (CT0-2, n=7 (cyan); CT12-14, n=7 (red)) or AAV-DIO-Clock-DN-P2A-EYFP (**K**) (CT0-2, n=6 (cyan-green); CT12-14, n=5 (pink)) into the adLA; unpaired t-test, two-tailed. **(L)** Representative confocal images of Per2 immunostaining in the adLA region at CT1 vs CT13 in WT mice in the presence (right) or absence (left) of pulsed TMT exposure across 4 days. DAPI and merged channels are also shown. Scale bars denote 50 μm. **(M)** Relative levels of Per2 intensity (represented as a fold-change relative to CT1 in controls) at CT1 vs CT13 in WT mice in the presence (TMT, purple; CT1, n=22; CT13, n=23) or absence (Control, green; CT1, n=20; CT13, n=22) of TMT. Data represented as simplified boxplots showing median with 1^st^ and 3^rd^ quartile boxes. N represents number of sections, and 3-4 mice were used for each condition; Mann-Whitney U tests with Bonferroni correction.

What advantages are conferred by the control of rhythmic behaviors by local clocks, rather than by a single master pacemaker? One potential answer may be more flexible, modular regulation of these extra-SCN oscillators and their circuits by distinct tissue-relevant cues. For example, the LA is well-known to function in fear and anxiety processing^22,23^, and so we asked whether clock cycling in these cells is impacted by these stimuli. We examined whether TMT (2,5-dihydro-2,4,5-trimethylthiazoline) exposure affects Per2 cycling in the adLA. TMT is an innate fear-inducing odorant from fox feces, and repeated exposure to TMT has been shown to alter Per2 cycling in the BLA^31^. Thus, we utilized a similar paradigm and quantified anti-Per2 immunostaining specifically in the adLA in WT mice. In control mice exposed to water, Per2 levels were higher at CT1, compared to CT13. In contrast, this cycling was reversed in mice with repeated exposure to TMT at ZT0.5, with significantly higher levels observed at CT13, compared to CT1 (Figure 6L and 6M). In contrast, repeated TMT exposure did not affect Per2 cycling in the SCN (Figure S7D and S7E). Taken together, these data demonstrate that local clock function is required in adLA^mWAKE^ neurons to generate rhythms of sensory perception and internal states and also suggest that fear-related cues can alter clock rhythms specifically in the adLA.

### mWAKE acts in adLA neurons to promote rhythmic behavior

Because our previous work in *Drosophila* suggested that WAKE acts downstream of the core clock, and because we find that mWAKE levels cycle in the LA, we asked whether the local clock in adLA neurons acts via mWAKE to regulate behavioral rhythms. To address whether mWAKE expression in the LA is under local clock control, we performed Western blot analyses using LA tissue from *mWake^(Cre/V^*^5^*^)^* mice bilaterally injected with AAV-DIO-EYFP or AAV-DIO-Clock-DN-P2A-EYFP virus into the adLA (Table S3). We found that expressing Clock-DN in adLA^mWAKE^ neurons significantly reduced mWAKE expression, compared to controls (Figure S7F and S7G). Next, we examined whether mWAKE was required in adLA^mWAKE^ neurons for daily rhythms of mechanical sensitivity and anxiety. To address this possibility, we performed bilateral injection of AAV-Cre-EGFP virus or control AAV-EGFP virus into the LA of heteroallelic mice carrying both a floxed allele and a null allele of *mWake* (*mWake^(flox/-)^*) (Figure 7A, 7B and Table S3). To confirm knockdown of *mWake* following viral Cre injection, we performed RNAscope *in situ* hybridization, which demonstrated substantial reduction of *mWake* transcript, likely due to nonsense-mediated decay (Figure S7H and S7I). Conditional knockout of mWAKE in the adLA eliminated cycling of cutaneous mechanical sensitivity and anxiety-like behavior as assessed by the EPM test (but not the OF test), such that the phenotypes during subjective night resembled those seen during the subjective day (Figure 7D, 7F, 7H, S7J, and S7K). In contrast, *mWake^(flox/-)^*mice with bilateral injection of control AAV-EGFP virus in the adLA exhibited normal daily rhythms of these behaviors (Figure 7C, 7E, 7G, S7J, and S7K). Taken together, these data suggest that a local clock mechanism in the adLA acts via mWAKE to generate rhythmic control of sensory perception and anxiety.

**Figure 7.**
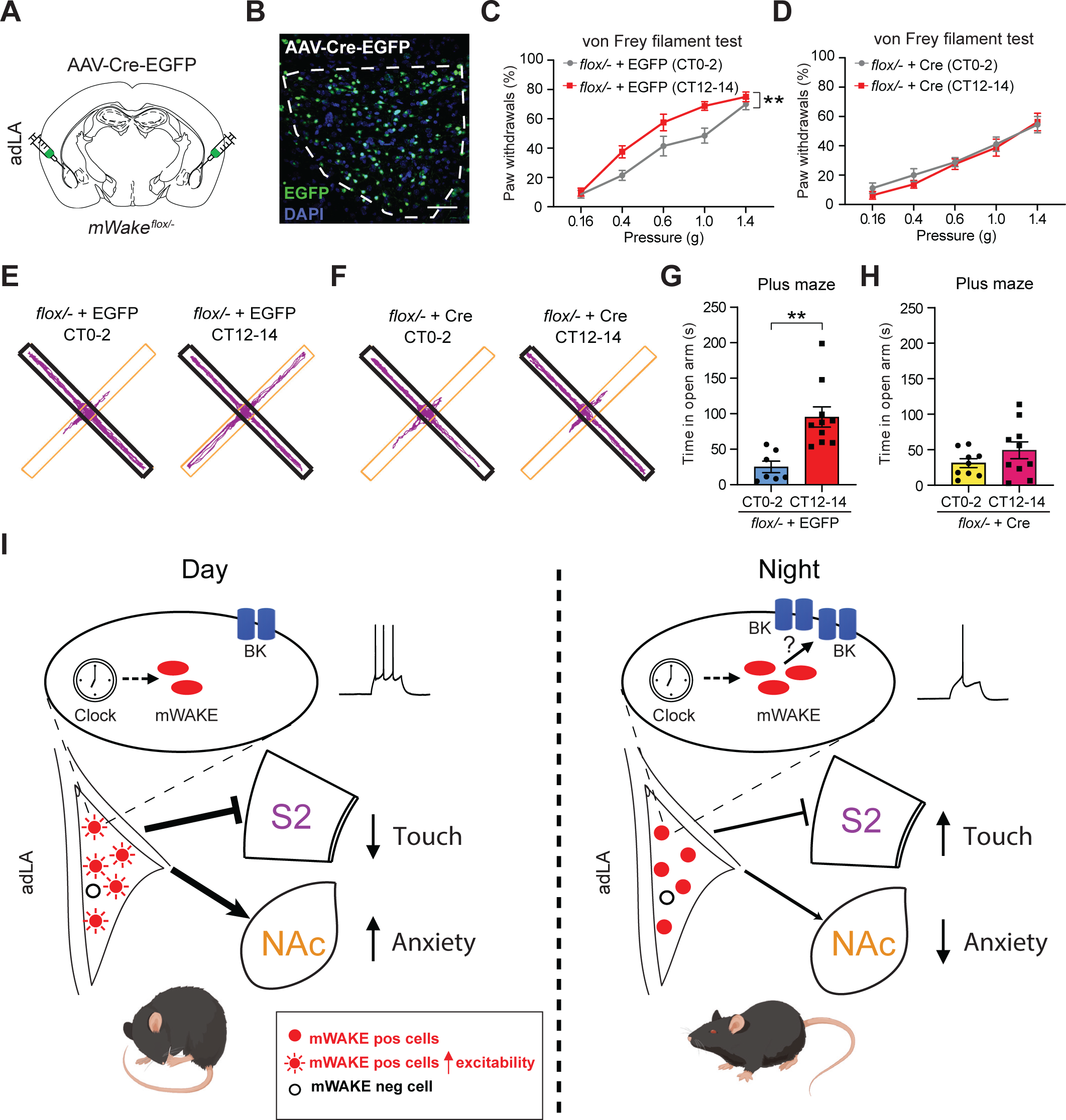
mWAKE mediates rhythmic control of mechanical sensitivity and anxiety. **(A)** Schematic showing bilateral injections of AAV-Cre-EGFP into the adLA of *mWake^(flox/-)^* mice. **(B)** Representative confocal image of the adLA for the mouse shown in (**A**), showing native GFP and DAPI signals. Scale bar denotes 50 μm. **(C** and **D)** Average hind foot withdrawal (%) in response to mechanical stimuli delivered using von Frey filaments of different strengths at CT0-2 (gray) or CT12-14 (red) for *mWake^(flox/-)^* mice injected with control AAV-EGFP (CT0-2, n=7; CT12-14, n=8) or AAV-Cre-EGFP (**D**) (CT0-2, n=9; CT12-14, n=8) into the adLA; two-way ANOVA with post-hoc Sidak. **(E** and **F)** Representative locomotor activity tracks for elevated plus maze assays at CT0-2 vs CT12-14 for *mWake^(flox/-)^* mice injected with control AAV-EGFP (**E**) (EGFP) or AAV-Cre-EGFP (**F**) (Cre) into the adLA. Closed and open arms indicated with thick black lines and thin yellow lines, respectively. **(G** and **H)** Average time spent in open arm during elevated plus maze assays at CT0-2 vs CT12-14 for *mWake^(flox/-)^* mice injected with control AAV-EGFP (**G**) (CT0-2, n=7 (cyan); CT12-14, n=10 (red)) or AAV-Cre-EGFP (**H**) (CT0-2, n=9 (yellow); CT12-14, n=10 (rose)) into the adLA; unpaired t-test, two-tailed. **(I)** Model. mWAKE expression defines a subset of LA neurons that function as an extra-SCN circadian oscillator. mWAKE levels rise at night under clock control. mWAKE, in turn, upregulates BK current at night, thus inhibiting the excitability of mWAKE-positive neurons (red circles), but not mWAKE-negative neurons (empty circles), at that time. This electrical cycling promotes greater activity of adLA^mWAKE^ neurons in the daytime and reduced activity of these cells in the nighttime. adLA^mWAKE^ neurons inhibit mechanical sensitivity and promote anxiety via projections to S2 and NAc, respectively. Thus, during the day, this circuit mechanism leads to a CLOCK- and mWAKE-dependent increase in anxiety (encouraging mice to seek shelter) and decrease in sensory perception (enhancing sleep amount or quality). Conversely, during their active period at night, these processes result in reduced anxiety (encouraging mice to explore) and enhanced sensory perception while engaging with the environment. Taken together with their function in regulating regional clock gene rhythms (Figure 2I), these findings suggest that adLAmWAKE neurons play a privileged role in organizing cellular and behavioral rhythms. Images of awake and sleeping mice were obtained from https://www.figdraw.com.

## DISCUSSION

By leveraging *mWak*e as a genetic marker, we identified a subpopulation of LA neurons that exhibits rhythmic intrinsic excitability and generates cycling of anxiety-like behaviors and mechanical sensitivity in a clock-dependent manner (Figure 7I). We show that mWAKE-positive LA neurons can tune anxiety-related behaviors via a non-canonical projection to the NAc, a region shown to be important for motivation, reward, and more recently, anxiety^36–38,40–42,48–50^. Interestingly, we also find that mWAKE-expressing neurons regulate touch and pain perception via a projection to S2. Traditionally, the LA has been considered the “sensory gateway” of the amygdala, and nearly all studies of the LA have focused on its role in integrating information from sensory cortex and transmitting this information to other subcortical structures^22,51–53^. However, it has long been recognized that LA neurons send prominent reciprocal projections to sensory-related neocortical areas, but the function of these projections is largely unknown^54,55^. Our findings show that LA^mWAKE^➔cortical projections can regulate sensory perception, and we speculate that this pathway provides a mechanism by which emotional state can modulate the perception of external stimuli.

In line with our previous studies in fly clock neurons and mouse DMH, we find that mWAKE suppresses intrinsic excitability of select LA neurons at night (Figure 7I). Selective knockout of mWAKE in adLA^mWAKE^ neurons not only abolishes circadian cycling of mechanical sensitivity and anxiety-like behaviors, but the resulting phenotypes (i.e., reduced mechanical sensitivity and increased anxiety) resemble those seen following gain-of-function manipulations of this circuit. A number of ion channels and pumps have been shown to mediate the oscillatory electrical activity of the SCN in mammals^56^. In contrast, the molecular mechanisms generating rhythms of neurophysiological activity in non-SCN brain regions are poorly understood^15,57^. In adLA^mWAKE^ neurons, we demonstrate that BK current cycles, with greater current amplitude at night and that this cycling depends on mWAKE.

What is the ethological relevance of the cycling of sensory perception and anxiety mediated by mWAKE and adLA^mWAKE^ circuits? During the day (the inactive period for mice), mWAKE levels are lower, resulting in reduced inhibition of adLA^mWAKE^ neurons. Increased activity of LA^mWAKE^ neurons, in turn, would increase anxiety, potentially encouraging the animals to seek shelter, and blunt sensory perception during sleep. Conversely, during the night (the active period for mice), mWAKE levels rise under clock control, decreasing adLA^mWAKE^ neuron activity. This reduced adLA^mwAKE^ signaling to NAc core and S2 would reduce anxiety to promote exploratory behavior and enhance sensory perception during the active period (Figure 7I).

Finally, multi-level hierarchical systems in the cerebral cortex are crucial for the generation of sensory-motor computations (e.g., Fuster’s hierarchy)^58^. Similarly, how behaviors are tuned by the internal representation of time is thought to depend on a network whereby the master pacemaker SCN coordinates the phase of local or peripheral oscillators, likely via secreted molecules and/or direct and indirect synaptic signaling^59–64^. However, although often described as a “hierarchy”, the SCN-dependent network has not been previously shown to contain intermediate level “nodes” that possess a privileged role in organizing molecular and behavioral rhythms across brain regions. We propose that adLA^mWAKE^ neurons function as an “intermediate”-level extra-SCN brain oscillator. Not only do these cells exhibit rhythmic clock gene expression and intrinsic excitability, they also regulate clock gene cycling in a different brain subregion. In addition, rather than that modulate a single behavior, “intermediate” oscillators, such as the adLA^mWAKE^ neurons, could organize “modules” of linked behaviors that are temporally coordinated in a daily manner. Finally, we find that Per2 rhythms in the LA are altered by exposure to TMT, a fear-inducing odor^31^, providing a potential mechanism by which the adLA^mWAKE^ extra-SCN oscillator could respond to relevant cues (e.g., fear or anxiety). Taken together, we hypothesize that adLA^mWAKE^ neurons amplify electrical or secreted rhythms from the SCN and coordinate this information across the LA and other brain regions. Because mWAKE is expressed in a number of brain regions that demonstrate clock gene cycling^20^, this molecule may broadly define intrinsically rhythmic extra-SCN oscillators that help coordinate circadian-related behaviors and information throughout the brain.

## Methods

### Animals

All animal procedures were approved by the Johns Hopkins Institutional Animal Care and Use Committee. All animals were group-housed unless otherwise specified and maintained with standard chow and water available *ad libitum*. Animals were raised in a common animal facility under a 14:10 hr Light:Dark (LD) cycle. Animals were further entrained to a 12:12 hr LD cycle in a satellite room for at least 2 weeks. Adult male mice were used for all experiments, unless otherwise specified. Genotyping was performed via Taq-Man based rtPCR probes (Transnetyx). Wild-type *C57BL/6J* mice were obtained from the Jackson Laboratory (JAX, 000664), and all mouse strains were backcrossed to this background at least seven times before use in behavioral experiments. The following previously described mouse strains were used in this study: an *mWake* null allele *(mWake^-^*)^29^, an *mWake* conditional allele (*mWake^flox^*)^29^, an *mWake* allele with a *tdTomato-P2A-Cre cassette* integrated into exon 5 (*mWake^Cre^*)^29^, and a tagged *mWake* allele (*mWake^V^*^5^)^20^. *VGLUT1^Cre^* mice were obtained from Jackson Laboratory (JAX, 037512).

### Molecular biology

The AAV-DIO-Clock-DN-P2A-EYFP and AAV-DO-Clock-DN-P2A-EYFP vectors and associated viruses were generated by Biohippo Inc. The Clock-DN sequence was based on a previously validated Clock-DN design utilized in *Drosophila*^32^ and synthesized with deletion of bases 91-141 followed by a P2A sequence. The Clock-DN-P2A sequence was inserted into an AAV-EF1α-DIO-EYFP or AAV-EF1α-DO-EYFP vector backbone using In-Fusion cloning (Takara Bio).

### *In situ* hybridization

RNAscope *in situ* hybridization was performed using the RNAscope® Multiplex Fluorescent Reagent Kit V2 (Advanced Cell Diagnostics/ACD, 323100) according to manufacturer’s instructions. Briefly, mice were anesthetized and then perfused with ice-cold 1xPBS (1.1 mM KH_2_PO_4_, 155.2 mM NaCl, 3.0 mM NaH_2_PO_4_, pH 7.4) and then 4% PFA in 1xPBS. Brains were removed and kept in 4% PFA for 24 hrs at 4°C. Following this, brains were immersed successively in 10%, 20%, and 30% sucrose in 1xPBS (for 1 day, 1 day, and 2 days, respectively at 4°C) until they sank to the bottom. Brains were then frozen in Tissue-Tek® Optimum Cutting Temperature (OCT) solution (VWR, 4583) with dry ice and stored in −80°C. 12-14 μm sections were obtained using a cryostat, placed on slides, and transferred to −80°C for storage. Sections were used for RNAscope *in situ* hybridization according to the ACDBio protocol (320293).

Target probes (mWake-C1 (1057631-C1), VGLUT2-C1 (319171-C1), VGAT-C2 (319191-C2), GRP-C2 (317861-C2), tdTomato-C3 (317041-C3)) were used in this paper. Opal^TM^ Dye (AKOYA Biosciences Opal^TM^ 4-Color Manual IHC Kit, NEL810001KT, 1:2000) was applied to develop the signal (520, 570, or 690). DAPI was then added to the sections and incubated for 30 sec at room temperature (RT). After removing the DAPI, ProLong™ Gold Antifade Mountant (Invitrogen, P36930) was immediately placed on each slide, followed by a glass coverslip.

Imaging was performed on a Zeiss LSM700 confocal microscope under 10x-40x magnification. For quantification of *mWake* RNAscope signal, average intensity was calculated using Fiji^65^. A maximum intensity projection was generated from confocal images of 14 μm thick-tissue slices containing the LA. The region of interest (ROI) was the LA and determined by the intensity and density of GFP signal from AAV-EGFP or AAV-Cre-EGFP.

### Immunostaining

Mice were anesthetized and perfused with ice-cold 1xPBS followed by 4% PFA in 1xPBS. After dissection, brains were kept in 4% PFA overnight. 40 µm coronal brain slices were collected using a vibratome (VT-1200s, Leica) and then washed twice with 1xPBS with 0.3% Triton X-100 (PBS-T). After blocking with PBS-T with 5% goat serum for 30 mins at RT, tissue sections were incubated with primary antibodies in PBS-T with 5% goat serum for 24 hrs at 4°C. The following primary antibodies were used: rabbit anti-Per2 (Millipore Sigma, AB2202, 1:500), mouse anti-CamKIIα (ThermoFisher Scientific, MA1-048, 1:200), mouse anti-SaCas9 (Epigentek, A-9001-100, 1:300), mouse anti-GAD67 (Abcam, ab26116, 1:200), or chicken anti-RFP (Rockland, 600-901-379, 1:1000). Sections were washed 4 times in PBS-T and then incubated with fluorescently labeled secondary antibodies in PBS-T with 5% goat serum overnight at 4°C. The following secondary antibodies were used: Alexa Fluor^TM^ 488 goat anti-rabbit (Invitrogen, A11008, 1:1000) for Per2 staining in WT mice, Alexa Fluor^TM^ 647 goat anti-rabbit (Invitrogen, A21244, 1:1000) for Per2 staining following viral injection, Alexa Fluor^TM^ 488 goat anti-mouse (Invitrogen, A11001, 1:1000) for CamKIIα, GAD67 or SaCas9 staining, and Alexa Fluor^TM^ 568 goat anti-chicken (Invitrogen, A11041, 1:1000) for tdTomato staining. After washing 4 times with PBS-T, mounting medium with DAPI (Vector Labs, H-1500-10) or Prolong Glass mounting media (Invitrogen, P36984) and coverslips were added. Imaging was performed on a Zeiss LSM700 confocal microscope under 10x-40x magnification.

For quantification of Per2 immunostaining in the LA, signals were analyzed using Imaris (v9.5.0, Oxford Instruments). ROIs were selected using DAPI or EGFP signal in Surface mode. After masking Per2 signal outside the ROIs, we adjusted the size of spots and the filters to detect signals in Spot mode, and then manually varied the number and location of spots until all Per2 signals in the ROI were detected. Per2 signal intensity in an individual neuron was quantified as the average intensity within a spot.

For analysis of projection patterns using ChR2 expression, 2-3 mo old male *mWake^(Cre/+)^* mice were injected with AAV-DIO-ChR2-EYFP virus in the adLA or pvLA (Table S3). After 2 wks for viral expression, anti-GFP immunostaining (rabbit anti-GFP, Invitrogen, A-11122, 1:500) on 60 mm coronal brain sections was performed as described above. Fluorescence images were taken using a Keyence microscope BZ-X710. Projection patterns from anterior forebrain (AP +2.33 mm from bregma) to midbrain (AP −5.91 mm from bregma) were identified according to the Paxinos and Franklin mouse brain atlas^66^. Target brain regions were subjectively scored from “+” to “++++” based on signal intensity.

### Single-cell RNA sequencing

#### sc/snRNA-Seq cell preparation

The lateral amygdalae were collected from 7-8 wk old male *mWake^(Cre/+)^* mice at ZT0-2 using a previously published protocol^67,68^. Briefly, the coronal plane (350 μm) containing the LA was sliced using a vibratome in bubbling aCSF. LA were dissected into Hibernate-A media with a 2% B-27 and GlutaMAX supplement (0.5 mM final concentration). Both LA from 2 mice were used for scRNA-Seq analysis, which was repeated with 2 additional mice. This procedure was also performed for snRNA-Seq analysis (in total, 8 mice were used).

For scRNA-Seq, tissues were dissociated in papain (Worthington) and debris was removed using OptiPrep density gradient media following cell dissociation. Cells were then processed immediately for scRNA-Seq.

For snRNA-Seq, tissues were processed following a modified 10x Genomics protocol. Lysis buffer containing (10 mM Tris-HCl pH 7.4, 10 mM NaCl, 3 mM MgCl_2_, 0.1% Tween-20, 0.1% Nonidet-P40, 0.01% Digitonin, 1 U/ml RNase inhibitor, 1% BSA) was added into a tube containing the lateral amygdalae and incubated on ice for 15 min with gentle pestle grinding (5 strokes every 3 min). Wash buffer (10 mM Tris-HCl pH 7.4, 10 mM NaCl, 3 mM MgCl_2_, 0.1% Tween-20, 0.2 U/ml RNase inhibitor, 1% BSA) was added, and nuclei were filtered through a 50 μm filter. Debris was removed using OptiPrep density gradient media and nuclei morphology was accessed under the light microscope. Nuclei were then immediately processed for snRNA-Seq.

#### scRNA-Seq/snRNA-Seq generation

Cells or nuclei from the lateral amygdalae were loaded into the 10x Genomics Chromium Single Cell System (10x Genomics) and libraries were generated using v3.1 chemistry following the manufacturer’s instructions. Two biological replicates for each scRNA-Seq and snRNA-Seq were conducted. Two technical replicates were used for one of the biological replicates.

Libraries were sequenced on an Illumina NovaSeq6000. scRNA-Seq data were first processed through Cell Ranger (v.5.0.0, 10x Genomics) with default parameters, aligned to the custom mm10 genome with *mWake* sequence^20^. snRNA-Seq data were processed with the same pipeline with ‘include-introns’. Matrix files were used for subsequent analysis.

#### scRNA-Seq/snRNA-Seq data analysis

Seurat v3.4^69^ was used to process the matrix files as previously described^70^. Cells with more than 500 genes, 2000 UMI, and less than 50% mitochondrial genes and 20% ribosomal genes were selected. Seurat ‘*scTransform*’ function was used to normalize the datasets by regressing the number of genes and UMIs using ‘*vars.to.regress*’. Batch variation was adjusted by Harmony v1.0^71^, by treating individual sc/snRNA-Seq run as a variance group, and UMAP dimensional reduction with Louvain clustering was used to identify cell types. scRNA-Seq and snRNA-Seq datasets were then merged with the integration function in Seurat. Neuronal clusters were subsetted and annotated into ‘adLA’, ‘pvLA’, and ‘BA’ which consists of glutamatergic neurons, and ‘Pericapsular region’ which is composed of GABAergic neurons. Annotations were based on a previous study^28^. Cells that display *mWake-Cre* and *Ankfn1* were labeled as ‘mWake-positive’ neurons and the rest were labeled as ‘mWake-negative’. Seurat ‘*FindMarkers*’ function was used to calculate differential expression between ‘mWake-positive’ and ‘mWake-negative’ neurons in ‘adLA’ or ‘pvLA’.

### Western blotting

LA tissue was dissected from *mWake^(V^*^5^*^/V^*^5^*^)^* or *mWake^(Cre/V^*^5^*^)^* male mice and then homogenized in RIPA solution (Millipore Sigma, R0278) containing 1:100 protease inhibitor cocktail (Millipore Sigma, P8340). After incubating on ice for 1 hr, lysates were cleared by centrifugation at 12,000g for 10 min. Protein concentration was measured using a Pierce BCA protein assay kit (ThermoFisher Scientific, 23227), and 20 μg protein was used to run on SDS-PAGE, followed by transfer to a PVDF membrane for 1.5 hrs at 100V. Membranes were blocked for 1 hr with Intercept ® Blocking Buffer (LI-COR, 927-60001) and then incubated overnight at 4°C with rabbit V5-Tag (D3H8Q) antibody (Cell Signaling Technology, 13202, 1:2000) in blocking buffer with 0.2% Tween 20. Following 4 washes with 1xTBST (137 mM NaCl, 20 mM Tris, 0.1% Tween 20, pH 7.6), blots were incubated with mouse anti-β-actin antibodies (Millipore Sigma, A5441, 1:5000) in Intercept ® Blocking Buffer for 1 hr at RT. Blots were again washed 4 times with 1xTBST and then incubated with anti-mouse (LI-COR, IRDye® 680RD, 926-68070) and anti-rabbit (LI-COR, IRDye® 800CW, 926-32211) fluorescent secondary antibodies (1:10,000) for 1 hr at RT. After 4 additional washes with 1xTBST, membranes were rinsed with 1xTBS and imaged on an Odyssey® Fc Imaging System (LI-COR Biosciences).

### Stereotaxic surgeries

8-10 wk old male mice were deeply anesthetized with a ketamine/xylazine mixture (100 mg/kg and 10 mg/kg). Mice were secured into a Stoelting stereotaxis frame, and all coordinates were zeroed when the glass pipette tip was placed on the bregma. Small (∼0.5 mm) craniotomies were performed to allow for virus injection (100-200 nl at ∼100 nl/min). Coordinates, volumes, viruses used, and their sources are listed in Table S3. Following injection, animals were allowed to recover and express viral genes for 2-5 weeks. Locations of viral injections were confirmed by post-hoc fluorescence imaging using 40-60 µm coronal brain slices.

### Behavioral Analyses

#### Mechanical sensitivity assay

Cutaneous mechanical sensitivity assays were performed essentially as previously described^72^. Under dim red light for visualization, 3-4 mo old mice were placed on a small metal mesh table and immediately covered by an inverted 500 ml glass beaker and then by a larger red plastic container. After a 20-30 min period to allow for habituation, an experimenter blinded to the genotype of the animal used von Frey filaments to deliver mechanical stimuli of varying strengths (0.16, 0.4, 0.6, 1.0, 1.4 g) to the hindfoot at CT0-2 or CT12-14. The same foot was tested 10x for each filament strength, and the % of withdrawals was recorded. Animals were only used for a single trial.

#### Open field

Open field (OF) assays were performed using a square chamber measuring 40 cm (length) x 40 cm (width) x 30 cm (height) made from white plastic. The chamber was wiped with 70% EtOH prior to use and allowed to dry completely (∼3 min). A single 3-4 mo old mouse was gently placed in the corner of the chamber and allowed to move freely for 5 min. ANY-maze video-tracking software (Stoelting Company) was used to record and analyze the video, sampling at 15 Hz. The center square of the open field comprised 50% of the total area, and the mouse was considered in the center square only when ≥75% of the mouse’s entire body was in this area.

#### Elevated plus maze

Elevated plus maze (EPM) assays were performed as previously described^73^. The apparatus used for the elevated plus maze has a cross shape with two open arms (30 x 5 cm), two closed arms (30 x 5 cm), and a center platform (5 x 5 cm). The closed arms have high (15 cm) walls enclosing the arms, and the entire apparatus is 30 cm above the floor.

The maze was wiped with 70% ethanol prior to use and allowed to dry completely (∼5 min). An individual 3-4 mo old mouse was gently placed in the center of the maze and allowed to move freely for 5 min. ANY-maze video-tracking software was used to record and analyze the video, sampling at 15 Hz, and the mouse was considered in the open or closed arm only when ≥75% of the mouse’s entire body was in the arm.

#### Wheel running

Wheel running data were collected using the ClockLab system (CLOCKLAB^TM^, ver. 3.604, Actimetrics). Individual 2-4 mo old mice were first placed in a wheel-running cage, and after a 5-7 day habituation period in LD, baseline activity in DD was recorded for 9 days (“Pre”). Animals were then randomly divided into two groups, and either AAV-DIO-Clock-DN-P2A-EYFP or AAV-DIO-EYFP virus was injected bilaterally into the SCN of these animals.

Mice were allowed to recover for 3 weeks after surgery, and then individual mice were again placed in a wheel-running cage. After a 5-7 day habituation period in LD, activity in DD was recorded for 9 days in DD (“Post”). Data were analyzed using the ClockLab Analysis program (CLOCKLAB^TM^, ver 6.1.02, Actimetrics).

#### Real-time place preference/avoidance

Real-time place preference/avoidance assays were performed as previously described^74,75^. The mice (11 weeks to 15 weeks) were placed in a behavioral arena (40 x 40 x 30 cm) for 20 min, and one side of the chamber was assigned as the “stimulation” side. Individual mice were placed in the non-stimulation side at the onset of the experiment. 20 Hz optical stimulation (20% on for 1 s alternating with a 1 s period of no stimulation using a blue LED at 3-8 mW) was delivered when the mouse crossed to the stimulation side; this stimulation ceased when the animal returned to the “non-stimulation” side.

### TMT exposure

2,5-dihydro-2,4,5-trimethylthiazoline (TMT, BioSRQ), a component of fox feces, was used as an innate stimulus to trigger fear. 2-3 mo old mice were removed from their home cages and placed in a new cage under dark conditions. The mice were then exposed to TMT by placing a kimwipe with 15 μl of TMT or 15 μl of water on the metal cage top for 15 min. Following this exposure, mice were returned to their original cage. Mice were exposed to TMT at ZT0.5 on 4 successive days in this manner. Animals were then anesthetized and perfused at CT1 or CT13, and Per2 immunostaining was performed as described above.

### Optogenetics

AAV-DIO-ChR2-EYFP virus was injected into the LA of 8-9 wk old *mWake^(Cre/+)^* mice as described above and in Table S3. Optical fibers (250 µm O.D.) with LED head mount (Amuza Inc) were inserted towards the LA, S2, or NAc core regions. Dental acrylic was applied to affix the ferrule on the skull. Mice were allowed to recover in a singly housed cage for 3-4 weeks after the surgery. To perform wireless optogenetic manipulation during behavioral assays, the headmount was connected to a wireless receiver (TeleR-1-P, Amuza Inc), and blue light stimulation was produced by a Teleopto Remote Controller controlled by a Pulse Generator (STOmk-2, Amuza Inc). For cutaneous mechanical sensitivity assays, optogenetic stimulation was delivered for 2 s (20 Hz, 20% on, 3-8 mW) while administering the von Frey filament.

Assays using different filaments were separated by ∼10 s intervals. For open field, elevated plus maze and place preference/avoidance assays, optogenetic stimulation was delivered for 1 s (20 Hz, 20% on, 3-8 mW), alternating with a 1 s period of no stimulation, throughout the test. These behavioral assays associated with optogenetic stimulation were performed at CT12-14.

### Chemogenetics

AAV-DIO-hm4D(Gi)-EGFP virus was injected bilaterally into the LA of 8-9 wk old *mWake^(Cre/+)^* mice as described above and in Table S3. Animals were allowed to recover 3-4 weeks after the surgery, and mechanical sensitivity, EPM, and OF tests were performed at CT0-2 within the following 2 weeks. CNO (10 mg/kg) was administered by intraperitoneal injection 30 min before performing behavioral assays.

### Electrophysiological recordings

Male mice (6-9 wks old) were used for patch-clamp recordings. Mice were deeply anesthetized with isoflurane, and brains were quickly removed and dissected in oxygenated (95% O_2_/5% CO_2_) ice-cold slicing solution (2.5 mM KCl, 1.25 mM NaH_2_PO_4_, 2 mM MgSO_4_, 2.5 mM CaCl_2_, 248 mM sucrose, 26 mM NaHCO_3_, 10 mM glucose). Coronal brain slices (250 μm) were obtained using a Leica vibratome (VT1200S), after which slices were recovered in 28L ACSF solution (124 mM NaCl, 2.5 mM KCl, 1.25 mM NaH_2_PO_4_, 2 mM MgSO_4_, 2.5 mM CaCl_2_, 26 mM NaHCO_3_, 10 mM glucose, osmolarity 290-300) for 30 mins, then incubated in RT for 1 hr. Slices were then transferred to an upright microscope (BX51WI, Olympus) and continuously perfused using oxygenated ACSF. Glass electrodes (5-8 MΩ) were filled with an internal solution containing 130 mM K-gluconate, 5 mM NaCl, 10 mM Na_2_C_4_H_8_N_3_O_5_P, 1 mM MgCl_2_, 0.2 mM EGTA, 10 mM HEPES, 2 mM MgATP, 0.5 mM Na_2_GTP, pH 7.2-7.3 with KOH, osmolarity 303). Whole-cell patch clamp recordings were obtained using a Multiclamp 700B amplifier (Molecular Devices). Data were sampled at 20 kHz, low-pass filtered at 2 kHz, and digitized using a Digidata 1440A (Molecular Devices). mWAKE-positive and -negative cells were visualized using infrared differential interference contrast (IR-DIC) and distinguished by native fluorescence. 0.5% biocytin (wt/wt) was added to the internal solution to label the recorded cell. Slices were fixed post-experiment in 4% PFA overnight, incubated with Alexa488-conjugated streptavidin (Invitrogen, 1:3000) for 24 hrs at 4 C°, immunostained for tdTomato as described above, and then imaged on a Zeiss 700 confocal microscope.

To measure neuronal excitability, 50 μM picrotoxin, 10 μM NBQX, and 50 μM D-AP5 were included in the bath solution (ACSF) to block synaptic inputs. The following parameters were measured: (1) *f-I* curve (−40 pA to 150 pA, 10 pA increment, 500 ms duration); (2) rheobase (100 ms duration); (3) Rin (the input resistance, measured using a hyperpolarizing current (−40 pA, 500 ms)); (4) RMP (resting membrane potential, measured at I=0).

For electrophysiological recordings coupled with optogenetic activation, mice were injected with AAV-DIO-ChR2-EYFP at 8-9 wks old as described above and in Table S3. Blue light stimulation was provided through the microscope objective using a pE-300^white^ LED illuminator (CoolLED, 10% power). Recordings were performed 3-5 weeks post-surgery, and the following parameters were measured: (1) action potentials (5 Hz, 10 Hz, 20 Hz, and 50 Hz stimulation (5 ms blue light pulse) was applied for 2 s to record action potentials under current-clamp configuration); (2) oEPSCs (optogenetically induced EPSC, 1 Hz, 5 ms blue light pulse) were recorded before and 10 mins after perfusion with ACSF containing 10 μM NBQX, and 50 μM D-AP5.

BK current was measured in voltage-clamp mode using an internal solution containing 120 mM K-gluconate, 20 mM KCl, 2 mM MgCl_2_, 2 mM K_2_-ATP, 0.5 mM Na-GTP, 10 mM HEPES, and 0.5 mM EGTA, pH 7.3. 1 μM tetrodotoxin (TTX) and 2 mM 4-aminopyridine were added to the ACSF. BK currents were elicited from a holding potential of −90 mV, stepping from −110 to +90 mV for 30 ms with 20 mV increments. Baseline currents were measured 3-4 times, and then, following focal application of the BK antagonist paxilline (10 μM), currents were recorded another 3-4 times. BK currents were isolated by subtracting the average currents after paxilline application from the average of the baseline currents.

### Statistical analysis

Statistical analyses were performed using Prism 8 (Graphpad). For comparisons of two normally distributed groups, unpaired Student t-tests were used; if these comparisons were before and after treatment of the cells, paired t-tests were used instead. For comparisons of more than two normally distributed groups, one-way ANOVA with post-hoc Tukey or Dunnett test was used.

For multiple comparisons with 2 factors, 2-way ANOVAs were followed by post-hoc Sidak tests. For comparisons of two groups of non-normally distributed data, Mann-Whitney U tests were performed. Bonferonni corrections were used, if needed, to correct for multiple comparisons. For comparisons of multiple groups of non-normally distributed data, Kruskal-Wallis tests were performed with post-hoc Dunn’s tests.

## Acknowledgements

We thank the Department of Neuroscience machine shop for building behavioral equipment and the Multiphoton Imaging Core for the use of an Imaris workstation. We thank the Transcriptomics and Deep Sequencing Core for scRNA sequencing. We thank members of the Wu Lab for discussion, and M. Tabuchi and R. Li for technical advice. This work was supported by a Kavli NDI Distinguished Graduate Student Fellowship (J.X.) and NIH grants R01MH126676 (S.B.), R01NS112266 (A.L.), and R35NS122181 (M.N.W.).

## Author Contributions

Q.L. and M.N.W. conceived the project, with input from C.A., S.B., and A.L. Q.L. performed all mouse genetics, behavioral experiments, surgeries, immunostaining, electrophysiological and optogenetic experiments. D.W.K. performed sc/sn RNA-Seq analysis. J.X. performed quantification for immunostaining experiments. S.S.L performed projection pattern analyses. B.J.B. generated mice used for this study. Q.L. and M.N.W. wrote the manuscript, with feedback from all authors.

## Declaration of interests

The authors declare no competing interests.

**Figure S1. LA^mWAKE^ neurons are excitatory, related to Figure 1**.

**(A)** UMAP plots showing sc/snRNA-Seq (single cell and single nucleus sequencing) clusters of the LA based on O’Leary et al, 2020^28^ (left). UMAP plots showing scRNA-Seq clusters and *mWake^tdTomato^* expression (top right). UMAP plots showing snRNA-Seq clusters and *mWake^tdTomato^*expression (bottom right).

**(B)** Representative confocal image of the amygdala of a WT mouse following RNAscope *in situ* hybridization using *mWake* (red) probe and DAPI dye (blue). adLA, central amygdala (CeA), and basal amygdala (BA) regions are outlined with dashed lines. Scale bar represents 200 μm.

**(C)** Representative confocal images of the adLA of a *mWake^(Cre/+)^* mouse following RNAscope *in situ* hybridization using tdTomato (red), *VGLUT2* (white), *VGAT* (green) probes and DAPI dye (blue, top) or *tdTomato* (red) and *gastrin-releasing peptide* (*GRP*, green) probes and DAPI dye (blue, bottom). Scale bars represent 50 μm.

**(D)** Representative confocal images showing immunostaining of tdTomato and CaMKIIα (upper) or tdTomato and GAD67 (lower) in the adLA of a *mWake^(Cre/+)^*mouse. DAPI and merged channels are also shown. Arrowheads indicate representative cells that are both mWAKE-positive and CaMKIIα-positive. Scale bars represent 50 μm.

**(E)** Percentage of CaMKIIα- and GAD67-positive neurons co-expressing mWAKE as defined by tdTomato labeling in an *mWake^(Cre/+)^*mouse (n=10 slices from 3 mice).

**Figure S2. Validation of AAV-DIO-Clock-DN-P2A-EYFP virus, related to Figure 2**

**(A)** Schematic showing bilateral injection of AAV-DIO-Clock-DN-P2A-EYFP into the SCN of VGAT*^(Cre/Cre)^* mice.

**(B)** Representative confocal image of EYFP fluorescence for the mouse shown in (**A**). Dashed lines outline the SCN. Scale bar represents 100 μm.

**(C** and **D)** Wheel-running traces of *VGAT^(Cre/Cre)^* mice before (Pre, left) and after (Post, right) injection of control AAV-DIO-EYFP (Control) (**C**) or AAV-DIO-Clock-DN-P2A-EYFP (Clock-DN) (**D**) virus into the SCN. 3 days of L:D and 9 days of D:D data are shown, with yellow bars indicating 12 hr light periods. X-axis represents time in hrs.

**(A) (E)** Chi-squared periodograms for the mice described in (**C**) and (**D**).

**(F** and **G)** Average period length (**F**) and circadian rhythm amplitude (**G**) for *VGAT^(Cre/Cre)^* mice before (Pre) and after (Post) injection of AAV-DIO-EYFP (n=3) or AAV-DIO-Clock-DN-P2A-EYFP (n=3) into the SCN; two-way ANOVA with post-hoc Sidak.

**Figure S3. Additional Per2 expression data, related to Figure 2**.

**(A)** Schematic showing bilateral injections of AAV-DIO-Clock-DN-P2A-EYFP virus in adLA with Per2 immunostaining from the pvLA of *mWake^(Cre/+)^* mice. Gray regions indicate pvLA.

**(B)** Representative confocal images of native EYFP and tdTomato signal or Per2 immunostaining in the pvLA region at CT1 vs CT13 in *mWake^(Cre/+)^*mice with AAV-DIO-EYFP or AAV-DIO-Clock-DN-P2A-EYFP injected bilaterally into the adLA. Merged channels are also shown. Scale bars denote 50 μm.

**(C** and **D)** Relative levels of Per2 intensity in mWAKE-positive (red) and mWAKE-negative (cyan) cells in the pvLA region at CT1 vs CT13 in *mWake^(Cre/+)^* mice with AAV-DIO-EYFP (CT1, n=12; CT13, n=11) (**C**) or AAV-DIO-Clock-DN-P2A-EYFP (CT1, n=11; CT13, n=10)

**(A) (D)** injected bilaterally into the adLA. Data represented as simplified boxplots showing median with 1^st^ and 3^rd^ quartile boxes. n represents number of sections, and 3 mice were used for each condition. Data represented as a fold-change relative to the signal for mWAKE-positive cells under control CT1 condition; Mann-Whitney U tests with Bonferroni correction.

**(B) (E)** Schematic image showing bilateral injections of AAV-DO-Clock-DN-EYFP virus into the pvLA of an *mWake^(Cre/+)^*mouse.

**(C) (F)** Representative confocal images of native EYFP and tdTomato signal or Per2 immunostaining in the pvLA region at CT1 vs CT13 in *mWake^(Cre/+)^* mice with AAV-DO-Clock-DN-P2A-EYFP virus injected bilaterally into pvLA. Merged channels are also shown. Scale bars denote 50 μm.

**(D) (G)** Relative levels of Per2 intensity in mWAKE-positive (pink) and mWAKE-negative cells (gray) in the pvLA region at CT1 (n=10) vs CT13 (n=12) for the mice shown in (**E**). For mWAKE-negative neurons, only cells expressing EYFP were quantified. Data represented as simplified boxplots showing median with 1^st^ and 3^rd^ quartile boxes. n represents number of sections, and 3 mice were used for each condition. Data represented as a fold-change relative to the signal for mWAKE-positive cells under control CT1 condition; Mann-Whitney U tests with Bonferroni correction.

**(E) (H)** Schematic showing bilateral injections of AAV-FLEX-Cas9-sgVGLUT1-sgVGLUT2 virus into adLA with Per2 immunostaining from the pvLA of *mWake^(Cre/+)^* mice. Gray regions indicate pvLA.

**(I)** Representative confocal images of native EYFP and tdTomato signal or Per2 immunostaining in the pvLA region at CT1 vs CT13 in *mWake^(Cre/+)^* mice with either AAV-FLEX-Cas9-sgLacZ or AAV-FLEX-Cas9-sgVGLUT1-sgVGLUT2 virus injected bilaterally into adLA. Merged channels are also shown. Cas9 expression in the adLA is shown in Fig. 2F. Scale bars denote 50 μm.

**(J** and **K)** Relative levels of Per2 intensity in mWAKE-positive (yellow) and mWAKE-negative (purple) cells in the pvLA region at CT1 vs CT13 in *mWake^(Cre/+)^*mice with either AAV-FLEX-Cas9-sgLacZ (CT1, n=10; CT13, n=8) (**J**) or AAV-FLEX-Cas9-sgVGLUT1-sgVGLUT2 (CT1, n=9; CT13, n=8) (**K**) injected bilaterally into the adLA. Data represented as simplified boxplots showing median with 1^st^ and 3^rd^ quartile boxes. n represents number of sections, and 3 mice were used for each condition. Data represented as a fold-change relative to the signal for mWAKE-positive cells under control CT1 condition; Mann-Whitney U tests with Bonferroni correction.

**(A) (L)** Schematic (left) and representative confocal images (right) showing unilateral injection of AAV-DIO-ChR2-EYFP into the adLA of a *mWake^Cre/+^* mouse and projections in pvLA. Scale bars denote 200 μm.

**(B) (M)** Schematic (left) and representative confocal images (right) showing unilateral injection of AAV-DIO-ChR2-EYFP in the pvLA of a *VGLUT1^(Cre/+)^* mouse and lack of projections in adLA. Scale bars denote 200 μm.

**Figure S4. Additional electrophysiological data, related to Figure 3**

**(A)** Representative confocal images showing tdTomato immunostaining and biocytin labeling of mWAKE-positive (upper) and -negative (lower) cells in the adLA from *mWake^(Cre/+)^* mice. Merged channels are also shown. Scale bars represent 20 μm.

**(B)** Mean resting membrane potential (RMP) of adLA^mWAKE^ neurons in *mWake^(Cre/+)^*(cyan) and *mWake^(Cre/Cre)^*(red) mice at ZT0-2 and ZT12-14 (data are from the same cells as in Figure 3D-3F).

**(C)** Mean resting membrane potential mWAKE-negative cells in the adLA from *mWake^(Cre/+)^* mice at ZT0-3 and ZT12-15 (data are from the same cells as in Figure 3H-3J).

**Figure S5. Projection pattern analysis of LA^mWAKE^ neurons, related to Figure 4**.

**(A)** Representative output areas of mWAKE-positive adLA neurons. Presynaptic targets of Cre-expressing LA neurons were visualized using a Cre-dependent AAV-DIO-ChR2-EYFP. Scale bars represent 1 mm. Confocal images of anti-EYFP immunostaining of injection site and additional coronal brain sections are shown with distance from bregma on the anterior-posterior (AP) axis indicated. Abbreviations: A25, cingulate cortex layer 25; ACx, auditory cortex; AID, agranular insular cortex, dorsal; BA, basal amygdala; CeA, central amygdala; Cla, claustrum; CPu, caudate/putamen; DI/GI, dysgranular/granular insular cortex; EA, extension of the amygdala; LOT, nucleus of the lateral olfactory tract; NAc core, nucleus accumbens core; PB, parabrachial nucleus; Po, posterior thalamic nuclear group, RRF, retrorubral field; S2, secondary somatosensory cortex; SN, substantia nigra; TeA, temporal association cortex; VP, ventral pallidum.

**(B)** adLA^mWAKE^ efferent intensities in cortical, subcortical, and midbrain regions, subjectively grouped into 4 categories, from weakest “+” to strongest “++++”. Projections anterior (left) and posterior (right) from the injection site are shown.

**Figure S6. Additional electrophysiological and behavioral data related to Figures 4 and 5**.

**(A)** Representative membrane potential traces (left) and amplitude (right) of S2 neuron EPSCs induced by optogenetic activation of adLA^mWAKE^ neurons in the absence (ACSF) or presence (NBQX+D-AP5) of ionotropic glutamate receptor blockers NBQX and D-AP5 (n=6); paired t-test, two-tailed. Blue bar indicates 5 ms blue light pulse.

**(B)** Representative membrane potential traces (left) and amplitude (right) of NAc neuron EPSCs induced by optogenetic activation of adLA^mWAKE^ neurons in the absence (ACSF) or presence (NBQX+D-AP5) of ionotropic glutamate receptor blockers NBQX and D-AP5 (n=9); paired t-test, two-tailed. Blue bar indicates 5 ms blue light pulse.

**(C)** Average time spent in center zone in open field assay 5 min before (left) and 5 min after (right) optogenetic stimulation of *mWake^(Cre/+)^* mice injected with control AAV-DIO-EYP (NAc-EYFP, gray, n=5) or AAV-DIO-ChR2-EYFP (NAc-ChR2, red, n=4) into the adLA with optical fibers implanted in NAc; unpaired t-test, two-tailed. Data are from the same animals in Figure 5H and 5I.

**(D)** Average time spent in open arm in elevated plus maze assays 5 min before (left) and 5 min after (right) optogenetic stimulation of *mWake^(Cre/+)^* mice injected with control AAV-DIO-EYP (NAc-EYFP, gray, n=5) or AAV-DIO-ChR2-EYFP (NAc-ChR2, red, n=5) into the adLA with optical fibers implanted in NAc; unpaired t-test, two-tailed. Data are from the same animals in Figure 5J and 5K.

**(E** and **F)** Average time spent in the center zone during the open field test (**E**) or in the open arm in the elevated plus maze test (**F**) for *mWake^(Cre/+)^* mice with either control AAV-DIO-EGFP (n=7, n=7, gray) or AAV-DIO-hM4D(Gi) (n=6, n=5, red) injected into the adLA and administered 10 mg/kg CNO; unpaired t-test, two-tailed. Data are from the same animals as in Figure 4K-4M.

**Figure S7. Additional behavioral and expression data, related to Figure 6 and Figure 7**.

**(A)** Average hind foot withdrawal (%) in response to mechanical stimuli delivered using von Frey filaments of different strengths at CT0-2 (gray, n=7) or CT12-14 (red, n=7) for WT mice; two-way ANOVA with post-hoc Sidak.

**(B)** Average time spent in center zone in open field assay at CT0-2 (cyan, n=10) vs CT12-14 (red, n=7) for WT mice; unpaired t-test, two-tailed.

**(C)** Average time spent in open arm during elevated plus maze assays at CT0-2 (cyan, n=9) vs CT12-14 (red, n=8) for WT mice; unpaired t-test, two-tailed.

**(D** and **E)** Representative confocal images of Per2 immunostaining (**D**) and relative levels of Per2 intensity (**E**) (represented as fold-change relative to CT1 under control condition) in the SCN at CT1 and CT13 in wild-type mice with (TMT: CT1, n=18; CT13, n=26) (purple) or without (Control: CT1, n=16; CT13, n=21) (green) TMT exposure. DAPI and merged channels are also shown. Data in (**E**) represented as simplified boxplots showing median with 1^st^ and 3^rd^ quartile boxes. Scale bars denote 50 μm. n represents number of sections, and 3 mice were used for each condition; Mann-Whitney U tests with Bonferroni correction.

**(F** and **G)** Representative immunoblot (**F**) and relative levels of mWAKE-V5 normalized to a β-actin loading control (**G**) from Western blot analyses of LA tissue at ZT13 using anti-V5 antibodies from *mWake^(Cre/V^*^5^*^)^* mice injected with control AAV-DIO-EYFP (EYFP, n=4, cyan) or AAV-DIO-Clock-DN-P2A-EYFP (Clock-DN, n=4, red) virus; unpaired t-test, two-tailed.

**(H** and **I)** Representative confocal images (**H**) and average *mWake* RNAscope signal intensity (**I**) with *mWake* (red) probe and GFP immunostaining in the LA of *mWake^(flox/-)^* mice injected with AAV-EGFP (n=13) or AAV-Cre-EGFP (n=16) virus into the LA. Merged channel also shown. Note that EGFP localizes to the cytoplasm, whereas Cre-EGFP localizes to the nuclei, due to the presence of a strong NLS. Scale bar represents 50 μm. n represents number of slices from 3-4 mice; Mann-Whitney U test.

**(J** and **K)** Representative locomotor activity tracks (**J**) and average time spent in center zone (**K**) in open field assay at CT0-2 vs CT12-14 for *mWake^(flox/-)^* mice injected with AAV-EGFP (*flox/-* + EGFP, left) (CT0-2, n=7 (cyan); CT12-14, n=9 (red)) or AAV-Cre-EGFP (*flox/-* + Cre, right) (CT0-2, n=9 (yellow); CT12-14, n=9 (rose)) virus into the LA; unpaired t-test, two-tailed. Center zone is indicated by yellow square.

**Table S1. adLA Differential Gene Expression**

Differential gene expression as measured by scRNA-seq in adLA between mWAKE-positive and mWAKE-negative neurons.

**Table S2. pvLA Differential Gene Expression**

Differential gene expression as measured by scRNA-seq in pvLA between mWAKE-positive and mWAKE-negative neurons.

**Table S3. Stereotaxic coordinates and viruses injected**

Details related to stereotaxic viral injections.

## Notes

### Competing Interest Statement

The authors have declared no competing interest.

## References

1. Mohawk, J.A., Green, C.B., and Takahashi, J.S. (2012). Central and peripheral circadian clocks in mammals. Annu Rev Neurosci 35, 445–462.

2. Herzog, E.D. (2007). Neurons and networks in daily rhythms. Nat Rev Neurosci 8, 790–802. 10.1038/nrn2215.

3. Ukai, H., and Ueda, H.R. (2010). Systems biology of mammalian circadian clocks. Annu Rev Physiol 72, 579–603. 10.1146/annurev-physiol-073109-130051.

4. Weaver, D.R. (1998). The suprachiasmatic nucleus: a 25-year retrospective. J Biol Rhythms 13, 100–112. 10.1177/074873098128999952.

5. Begemann, K., Neumann, A.M., and Oster, H. (2020). Regulation and function of extra-SCN circadian oscillators in the brain. Acta Physiol (Oxf) 229, e13446. 10.1111/apha.13446.

6. Hasegawa, S., Fukushima, H., Hosoda, H., Serita, T., Ishikawa, R., Rokukawa, T., Kawahara-Miki, R., Zhang, Y., Ohta, M., Okada, S., et al. (2019). Hippocampal clock regulates memory retrieval via Dopamine and PKA-induced GluA1 phosphorylation. Nat Commun 10, 5766. 10.1038/s41467-019-13554-y.

7. Koronowski, K.B., and Sassone-Corsi, P. (2021). Communicating clocks shape circadian homeostasis. Science 371. 10.1126/science.abd0951.

8. Porcu, A., Vaughan, M., Nilsson, A., Arimoto, N., Lamia, K., and Welsh, D.K. (2020). Vulnerability to helpless behavior is regulated by the circadian clock component CRYPTOCHROME in the mouse nucleus accumbens. Proc Natl Acad Sci U S A 117, 13771–13782. 10.1073/pnas.2000258117.

9. Koronowski, K.B., Kinouchi, K., Welz, P.S., Smith, J.G., Zinna, V.M., Shi, J., Samad, M., Chen, S., Magnan, C.N., Kinchen, J.M., et al. (2019). Defining the Independence of the Liver Circadian Clock. Cell 177, 1448–1462 e1414. 10.1016/j.cell.2019.04.025.

10. Fuller, P.M., Lu, J., and Saper, C.B. (2008). Differential rescue of light-and food-entrainable circadian rhythms. Science 320, 1074–1077. 10.1126/science.1153277.

11. Mistlberger, R.E., Buijs, R.M., Challet, E., Escobar, C., Landry, G.J., Kalsbeek, A., Pevet, P., and Shibata, S. (2009). Food anticipation in Bmal1-/-and AAV-Bmal1 rescued mice: a reply to Fuller et al. J Circadian Rhythms 7, 11. 10.1186/1740-3391-7-11.

12. Abe, M., Herzog, E.D., Yamazaki, S., Straume, M., Tei, H., Sakaki, Y., Menaker, M., and Block, G.D. (2002). Circadian rhythms in isolated brain regions. J Neurosci 22, 350–356. 10.1523/JNEUROSCI.22-01-00350.2002.

13. Feillet, C.A., Mendoza, J., Albrecht, U., Pevet, P., and Challet, E. (2008). Forebrain oscillators ticking with different clock hands. Mol Cell Neurosci 37, 209–221. 10.1016/j.mcn.2007.09.010.

14. Guilding, C., and Piggins, H.D. (2007). Challenging the omnipotence of the suprachiasmatic timekeeper: are circadian oscillators present throughout the mammalian brain? Eur J Neurosci 25, 3195–3216. 10.1111/j.1460-9568.2007.05581.x.

15. Paul, J.R., Davis, J.A., Goode, L.K., Becker, B.K., Fusilier, A., Meador-Woodruff, A., and Gamble, K.L. (2020). Circadian regulation of membrane physiology in neural oscillators throughout the brain. Eur J Neurosci 51, 109–138. 10.1111/ejn.14343.

16. Lowrey, P.L., and Takahashi, J.S. (2011). Genetics of circadian rhythms in Mammalian model organisms. Adv Genet 74, 175–230.

17. Allada, R., Emery, P., Takahashi, J.S., and Rosbash, M. (2001). Stopping time: the genetics of fly and mouse circadian clocks. Annu Rev Neurosci 24, 1091–1119.

18. Liu, S., Lamaze, A., Liu, Q., Tabuchi, M., Yang, Y., Fowler, M., Bharadwaj, R., Zhang, J., Bedont, J., Blackshaw, S., et al. (2014). WIDE AWAKE mediates the circadian timing of sleep onset. Neuron 82, 151–166. 10.1016/j.neuron.2014.01.040.

19. Tabuchi, M., Monaco, J.D., Duan, G., Bell, B., Liu, S., Liu, Q., Zhang, K., and Wu, M.N. (2018). Clock-Generated Temporal Codes Determine Synaptic Plasticity to Control Sleep. Cell 175, 1213–1227 e1218. 10.1016/j.cell.2018.09.016.

20. Bell, B.J., Wang, A.A., Kim, D.W., Xiong, J., Blackshaw, S., and Wu, M.N. (2021). Characterization of mWake expression in the murine brain. J Comp Neurol 529, 1954–1987. 10.1002/cne.25066.

21. Zhang, S., Ross, K.D., Seidner, G.A., Gorman, M.R., Poon, T.H., Wang, X., Keithley, E.M., Lee, P.N., Martindale, M.Q., Joiner, W.J., and Hamilton, B.A. (2015). Nmf9 Encodes a Highly Conserved Protein Important to Neurological Function in Mice and Flies. PLoS Genet 11, e1005344. 10.1371/journal.pgen.1005344.

22. Janak, P.H., and Tye, K.M. (2015). From circuits to behaviour in the amygdala. Nature 517, 284–292. 10.1038/nature14188.

23. LeDoux, J.E. (2000). Emotion circuits in the brain. Annu Rev Neurosci 23, 155–184. 10.1146/annurev.neuro.23.1.155.

24. Yang, Y., and Wang, J.Z. (2017). From Structure to Behavior in Basolateral Amygdala-Hippocampus Circuits. Front Neural Circuits 11, 86. 10.3389/fncir.2017.00086.

25. Repa, J.C., Muller, J., Apergis, J., Desrochers, T.M., Zhou, Y., and LeDoux, J.E. (2001). Two different lateral amygdala cell populations contribute to the initiation and storage of memory. Nat Neurosci 4, 724–731. 10.1038/89512.

26. Johnson, L.R., Hou, M., Ponce-Alvarez, A., Gribelyuk, L.M., Alphs, H.H., Albert, L., Brown, B.L., Ledoux, J.E., and Doyere, V. (2008). A recurrent network in the lateral amygdala: a mechanism for coincidence detection. Front Neural Circuits 2, 3. 10.3389/neuro.04.003.2008.

27. Wilson, Y.M., and Murphy, M. (2009). A discrete population of neurons in the lateral amygdala is specifically activated by contextual fear conditioning. Learn Mem 16, 357–361. 10.1101/lm.1361509.

28. O’Leary, T.P., Sullivan, K.E., Wang, L., Clements, J., Lemire, A.L., and Cembrowski, M.S. (2020). Extensive and spatially variable within-cell-type heterogeneity across the basolateral amygdala. Elife 9. 10.7554/eLife.59003.

29. Liu, Q., Bell, B.J., Kim, D.W., Lee, S.S., Keles, M.F., Liu, Q., Blum, I.D., Wang, A.A., Blank, E.J., Xiong, J., Bedont, J.L., Chang, A.J., Issa, H., Cohen, J.Y., Blackshaw, S., Wu, M.N. (2023). A Clock-Dependent Brake for Rhythmic Arousal in the Dorsomedial Hypothalamus. Nat Commun, in press. 10.1038/s41467-023-41877-4.

30. Lamont, E.W., Robinson, B., Stewart, J., and Amir, S. (2005). The central and basolateral nuclei of the amygdala exhibit opposite diurnal rhythms of expression of the clock protein Period2. Proc Natl Acad Sci U S A 102, 4180–4184. 10.1073/pnas.0500901102.

31. Pantazopoulos, H., Dolatshad, H., and Davis, F.C. (2011). A fear-inducing odor alters PER2 and c-Fos expression in brain regions involved in fear memory. PLoS One 6, e20658. 10.1371/journal.pone.0020658.

32. Tanoue, S., Krishnan, P., Krishnan, B., Dryer, S.E., and Hardin, P.E. (2004). Circadian clocks in antennal neurons are necessary and sufficient for olfaction rhythms in Drosophila. Curr Biol 14, 638–649. 10.1016/j.cub.2004.04.009.

33. Hannibal, J. (2002). Neurotransmitters of the retino-hypothalamic tract. Cell Tissue Res 309, 73–88. 10.1007/s00441-002-0574-3.

34. Meijer, J.H., and Schwartz, W.J. (2003). In search of the pathways for light-induced pacemaker resetting in the suprachiasmatic nucleus. J Biol Rhythms 18, 235–249. 10.1177/0748730403018003006.

35. Hunker, A.C., Soden, M.E., Krayushkina, D., Heymann, G., Awatramani, R., and Zweifel, L.S. (2020). Conditional Single Vector CRISPR/SaCas9 Viruses for Efficient Mutagenesis in the Adult Mouse Nervous System. Cell Rep 30, 4303–4316 e4306. 10.1016/j.celrep.2020.02.092.

36. Kalivas, P.W., and Nakamura, M. (1999). Neural systems for behavioral activation and reward. Curr Opin Neurobiol 9, 223–227. 10.1016/s0959-4388(99)80031-2.

37. Klawonn, A.M., and Malenka, R.C. (2018). Nucleus Accumbens Modulation in Reward and Aversion. Cold Spring Harb Symp Quant Biol 83, 119–129. 10.1101/sqb.2018.83.037457.

38. Gale, J.T., Shields, D.C., Ishizawa, Y., and Eskandar, E.N. (2014). Reward and reinforcement activity in the nucleus accumbens during learning. Front Behav Neurosci 8, 114. 10.3389/fnbeh.2014.00114.

39. Levita, L., Hoskin, R., and Champi, S. (2012). Avoidance of harm and anxiety: a role for the nucleus accumbens. Neuroimage 62, 189–198. 10.1016/j.neuroimage.2012.04.059.

40. Gunaydin, L.A., and Kreitzer, A.C. (2016). Cortico-Basal Ganglia Circuit Function in Psychiatric Disease. Annu Rev Physiol 78, 327–350. 10.1146/annurev-physiol-021115-105355.

41. Gebara, E., Zanoletti, O., Ghosal, S., Grosse, J., Schneider, B.L., Knott, G., Astori, S., and Sandi, C. (2021). Mitofusin-2 in the Nucleus Accumbens Regulates Anxiety and Depression-like Behaviors Through Mitochondrial and Neuronal Actions. Biol Psychiatry 89, 1033–1044. 10.1016/j.biopsych.2020.12.003.

42. Zhang, X.Y., Peng, S.Y., Shen, L.P., Zhuang, Q.X., Li, B., Xie, S.T., Li, Q.X., Shi, M.R., Ma, T.Y., Zhang, Q., et al. (2020). Targeting presynaptic H3 heteroreceptor in nucleus accumbens to improve anxiety and obsessive-compulsive-like behaviors. Proc Natl Acad Sci U S A 117, 32155–32164. 10.1073/pnas.2008456117.

43. Daguet, I., Raverot, V., Bouhassira, D., and Gronfier, C. (2022). Circadian rhythmicity of pain sensitivity in humans. Brain 145, 3225–3235. 10.1093/brain/awac147.

44. Griebel, G., Moreau, J.L., Jenck, F., Martin, J.R., and Misslin, R. (1993). Some critical determinants of the behaviour of rats in the elevated plus-maze. Behav Processes 29, 37–47. 10.1016/0376-6357(93)90026-N.

45. Bertoglio, L.J., and Carobrez, A.P. (2002). Behavioral profile of rats submitted to session 1-session 2 in the elevated plus-maze during diurnal/nocturnal phases and under different illumination conditions. Behav Brain Res 132, 135–143. 10.1016/s0166-4328(01)00396-5.

46. Andrade, M.M., Tome, M.F., Santiago, E.S., Lucia-Santos, A., and de Andrade, T.G. (2003). Longitudinal study of daily variation of rats’ behavior in the elevated plus-maze. Physiol Behav 78, 125–133. 10.1016/s0031-9384(02)00941-1.

47. Yannielli, P.C., Kanterewicz, B.I., and Cardinali, D.P. (1996). Circadian Changes in Anxiolysis-Related Behavior of Syrian Hamsters. Correlation with Hypothalamic GABA Release. Biological Rhythm Research 27, 365–373.

48. Day, J.J., and Carelli, R.M. (2007). The nucleus accumbens and Pavlovian reward learning. Neuroscientist 13, 148–159. 10.1177/1073858406295854.

49. Salgado, S., and Kaplitt, M.G. (2015). The Nucleus Accumbens: A Comprehensive Review. Stereotact Funct Neurosurg 93, 75–93. 10.1159/000368279.

50. Castro, D.C., and Bruchas, M.R. (2019). A Motivational and Neuropeptidergic Hub: Anatomical and Functional Diversity within the Nucleus Accumbens Shell. Neuron 102, 529–552. 10.1016/j.neuron.2019.03.003.

51. McDonald, A.J. (1998). Cortical pathways to the mammalian amygdala. Prog Neurobiol 55, 257–332. 10.1016/s0301-0082(98)00003-3.

52. Tovote, P., Fadok, J.P., and Luthi, A. (2015). Neuronal circuits for fear and anxiety. Nat Rev Neurosci 16, 317–331. 10.1038/nrn3945.

53. LeDoux, J. (2007). The amygdala. Curr Biol 17, R868–874. 10.1016/j.cub.2007.08.005.

54. Price, J.L., Russchen, F.T., and Amaral, D.G. (1987). The limbic region II: The amygdala complex. In Handbook of Chemical Neuroanatomy, Vol. 5, 279-388. Eds. A. Bjorklund, T. Hokfelt, and L.W. Swanson. Elsevier: Amsterdam.

55. Pitkanen, A. (2000). Connectivity of the rat amygdaloid complex. in The Amygdala: A Functional Analysis (2nd ed.) 31-115. Ed. J. Aggleton. Oxford University Press: NY.

56. Colwell, C.S. (2011). Linking neural activity and molecular oscillations in the SCN. Nat Rev Neurosci 12, 553–569. 10.1038/nrn3086.

57. Gonzalez, J.C., Lee, H., Vincent, A.M., Hill, A.L., Goode, L.K., King, G.D., Gamble, K.L., Wadiche, J.I., and Overstreet-Wadiche, L. (2023). Circadian regulation of dentate gyrus excitability mediated by G-protein signaling. Cell Rep 42, 112039. 10.1016/j.celrep.2023.112039.

58. Botvinick, M.M. (2007). Multilevel structure in behaviour and in the brain: a model of Fuster’s hierarchy. Philos Trans R Soc Lond B Biol Sci 362, 1615–1626. 10.1098/rstb.2007.2056.

59. Cheng, M.Y., Bullock, C.M., Li, C.Y., Lee, A.G., Bermak, J.C., Belluzzi, J., Weaver, D.R., Leslie, F.M., and Zhou, Q.Y. (2002). Prokineticin 2 transmits the behavioural circadian rhythm of the suprachiasmatic nucleus. Nature 417, 405–410. Doi 10.1038/417405a.

60. Kramer, A., Yang, F.C., Snodgrass, P., Li, X., Scammell, T.E., Davis, F.C., and Weitz, C.J. (2001). Regulation of daily locomotor activity and sleep by hypothalamic EGF receptor signaling. Science 294, 2511–2515.

61. Pevet, P., and Challet, E. (2011). Melatonin: both master clock output and internal time-giver in the circadian clocks network. J Physiol Paris 105, 170–182. 10.1016/j.jphysparis.2011.07.001.

62. Aston-Jones, G., Chen, S., Zhu, Y., and Oshinsky, M.L. (2001). A neural circuit for circadian regulation of arousal. Nat Neurosci 4, 732–738. 10.1038/89522.

63. Chou, T.C., Scammell, T.E., Gooley, J.J., Gaus, S.E., Saper, C.B., and Lu, J. (2003). Critical role of dorsomedial hypothalamic nucleus in a wide range of behavioral circadian rhythms. J Neurosci 23, 10691–10702.

64. Mistlberger, R.E. (2005). Circadian regulation of sleep in mammals: role of the suprachiasmatic nucleus. Brain Res Brain Res Rev 49, 429–454.

65. Schindelin, J., Arganda-Carreras, I., Frise, E., Kaynig, V., Longair, M., Pietzsch, T., Preibisch, S., Rueden, C., Saalfeld, S., Schmid, B., et al. (2012). Fiji: an open-source platform for biological-image analysis. Nature methods 9, 676-682. 10.1038/nmeth.2019.

66. Paxinos, G., and Franklin, K. (2012). The Mouse Brain in Stereotaxic Coordinates (4th Edition). Academic Press: San Diego.

67. Kim, D.W., Liu, K., Wang, Z.Q., Zhang, Y.S., Bathini, A., Brown, M.P., Lin, S.H., Washington, P.W., Sun, C., Lindtner, S., et al. (2021). Gene regulatory networks controlling differentiation, survival, and diversification of hypothalamic Lhx6-expressing GABAergic neurons. Commun Biol 4, 95. 10.1038/s42003-020-01616-7.

68. Kim, D.W., Washington, P.W., Wang, Z.Q., Lin, S.H., Sun, C., Ismail, B.T., Wang, H., Jiang, L., and Blackshaw, S. (2020). The cellular and molecular landscape of hypothalamic patterning and differentiation from embryonic to late postnatal development. Nat Commun 11, 4360. 10.1038/s41467-020-18231-z.

69. Stuart, T., Butler, A., Hoffman, P., Hafemeister, C., Papalexi, E., Mauck, W.M., 3rd, Hao, Y., Stoeckius, M., Smibert, P., and Satija, R. (2019). Comprehensive Integration of Single-Cell Data. Cell 177, 1888-1902 e1821. 10.1016/j.cell.2019.05.031.

70. Kim, D.W., Tu, K.J., Wei, A., Lau, A.J., Gonzalez-Gil, A., Cao, T., Braunstein, K., Ling, J.P., Troncoso, J.C., Wong, P.C., et al. (2022). Amyloid-beta and tau pathologies act synergistically to induce novel disease stage-specific microglia subtypes. Mol Neurodegener 17, 83. 10.1186/s13024-022-00589-x.

71. Korsunsky, I., Millard, N., Fan, J., Slowikowski, K., Zhang, F., Wei, K., Baglaenko, Y., Brenner, M., Loh, P.R., and Raychaudhuri, S. (2019). Fast, sensitive and accurate integration of single-cell data with Harmony. Nat Methods 16, 1289–1296. 10.1038/s41592-019-0619-0.

72. Liu, Y., Latremoliere, A., Li, X., Zhang, Z., Chen, M., Wang, X., Fang, C., Zhu, J., Alexandre, C., Gao, Z., et al. (2018). Touch and tactile neuropathic pain sensitivity are set by corticospinal projections. Nature 561, 547–550. 10.1038/s41586-018-0515-2.

73. Smith, G.W., Aubry, J.M., Dellu, F., Contarino, A., Bilezikjian, L.M., Gold, L.H., Chen, R., Marchuk, Y., Hauser, C., Bentley, C.A., et al. (1998). Corticotropin releasing factor receptor 1-deficient mice display decreased anxiety, impaired stress response, and aberrant neuroendocrine development. Neuron 20, 1093–1102. 10.1016/s0896-6273(00)80491-2.

74. Zhang, Z., Liu, Q., Wen, P., Zhang, J., Rao, X., Zhou, Z., Zhang, H., He, X., Li, J., Zhou, Z., et al. (2017). Activation of the dopaminergic pathway from VTA to the medial olfactory tubercle generates odor-preference and reward. Elife 6. 10.7554/eLife.25423.

75. Do-Monte, F.H., Minier-Toribio, A., Quinones-Laracuente, K., Medina-Colon, E.M., and Quirk, G.J. (2017). Thalamic Regulation of Sucrose Seeking during Unexpected Reward Omission. Neuron 94, 388–400 e384. 10.1016/j.neuron.2017.03.036.

