## Supplemental Table 3 for "An Amygdalar Oscillator Coordinates Cellular and Behavioral Rhythms"

**Table S3. Stereotaxic coordinates and viruses injected**

| Location | Coordinates | Virus (Source) | Use | Vol. | Laterality | |
| --- | --- | --- | --- | --- | --- | --- |
| adLA | AP:-0.95, ML:+3.40, DV:-4.07 | AAV1-pCAG-FLEX-EGFP-WPRE (Addgene# 51502-AAV1)  “AAC-FLEX-EGFP” | Projection  Mapping | 100nl | | Uni- |
|  | AP:-0.95, ML:+3.40, DV:-4.07 | AAV-EF1α-double floxed-hChR2(H134R)-EYFP (Addgene# 20298-AAV9) “AAV-DIO-ChR2” | Projection mapping | 100nl | | Uni- |
|  | AP:-0.95, ML:+/-3.40, DV:-4.07 | AAV9-Ef1α-DIO-EYFP (Addgene# 27056-AAV9) “AAV-DIO-EYFP” | Control | 150nl | | Bi- |
|  | AP:-0.95, ML:+/-3.40, DV:-4.07 | AAV-EF1α-double floxed-hChR2(H134R)-EYFP (Addgene# 20298-AAV9) “AAV-DIO-ChR2” | Optogenetic activation | 150nl | | Bi- |
|  | AP:-0.95, ML:+/-3.40, DV:-4.07 | AAV1-pCAG-FLEX-EGFP-WPRE  (Addgene# 51502-AAV1) “AAC-FLEX-EGFP” | Control | 150nl | | Bi |
|  | AP:-0.95, ML:+/-3.40, DV:-4.07 | AAV2/9-Ef1α-DIO-hM4D(Gi)-EGFP-WPREs  (BioHippo)  “AAV-DIO-hM4D(Gi)” | Dread inhibition | 150nl | | Bi- |
|  | AP:-0.95, ML:+/-3.40, DV:-4.07 | AAV2/9-EF1α-DIO-Clock-DN-P2A-EYFP (BioHippo)  “AAV-DIO-Clock-DN-P2A-EYFP” | Cre-on  Clock-DN | 150nl | | Bi- |
|  | AP:-0.95, ML:+/-3.40, DV:-4.07 | AAV9-hSyn-EGFP  (Addgene# 50465-AAV9)  “AAV-EGFP” | Control | 100nl | | Bi- |
|  | AP:-0.95, ML:+/-3.40, DV:-4.07 | AAV9-hSyn-HI-EGFP-Cre-WPRE-SV40  (Addgene# 105540-AAV9)  “AAV-Cre-EGFP” | Conditional  knockout | 100nl | | Bi- |
|  | AP:-0.95, ML:+/-3.40, DV:-4.07 | AAV2/9-EF1α-FLEX-SaCas9-U6-sgLacZ  “AAV-FLEX-sgLacZ” (BioHippo) | control | 150nl | | Bi- |
|  | AP:-0.95, ML:+/-3.40, DV:-4.07 | AAV2/9-CMV-FLEX-SaCas9-U6-sgVglut1-P2A-U6-sgVGlut2  “AAV-FLEX-sgVGLUT1-sgVGLUT2” (BioHippo) | GLUT knockdown | 150nl | | Bi- |
| pvLA | AP:-1.95  ML:+/-3.52  DV:-4.40 | AAV2/9-EF1α-DO-Clock-DN-P2A-EYFP (BioHippo)  “AAV-DO-Clock-DN-P2A-EYFP” | Cre-off  Clock-DN | 150nl | | Bi- |
|  | AP:-1.95  ML:+/-3.52  DV:-4.40 | AAV-EF1α-double floxed-hChR2(H134R)-EYFP (Addgene# 20298-AAV9)  “AAV-DIO-ChR2” | Projection  Mapping | 150nl | | Uni- |
| SCN | AP:-0.4, ML:+/-0.18, DV:-5.75 | AAV9-Ef1α-DIO-EYFP (Addgene# 27056-AAV9) “AAV-DIO-EYFP” | control | 150nl | | Bi- |
|  | AP:-0.4, ML:+/-0.18, DV:-5.75 | AAV2/9-EF1α-DIO-Clock-DN-EYFP (BioHippo)  “AAV-DIO-Clock-DN-EYFP” | Cre-on  Clock-DN | 150nl | | Bi- |
| adLA | AP:-0.95, ML:+/-3.40, DV:-4.02 |  | Optic fiber  implantation |  | | Bi- |
| S2 | AP:+0.13, ML:+/-3.50, DV:-3.2 |  | Optic fiber  implantation |  | | Bi- |
| NAc core | AP:+1.25, ML:+/-1.4, DV:-4.5 |  | Optic fiber  implantation |  | | Bi- |
